# Comparative analysis of naked mole-rat thermogenesis and its potential to maintain euthermia in response to cold

**DOI:** 10.1101/2025.07.11.664220

**Authors:** Aleksei Mikhalchenko, June K. Corrigan, Yuchen He, Zalan Peterfi, Sun Hee Yim, Sang-Goo Lee, Zhaoming Deng, Vince G. Amoroso, Vera Gorbunova, Andrei Seluanov, Thomas J. Park, Alexander S. Banks, Vadim N. Gladyshev

**Affiliations:** Division of Genetics, Department of Medicine, Brigham and Women’s Hospital, Harvard Medical School, Boston, MA 02115, USA; Division of Endocrinology, Diabetes and Metabolism, Beth Israel Deaconess Medical Center, Harvard Medical School, Boston, MA 02215, USA; Department of Biological Sciences, University of Illinois at Chicago, Chicago, IL 60607, USA; University of Rochester, Department of Biology, Rochester, NY 14627, USA

**Keywords:** naked mole-rat, thermogenesis, UCP1, brown adipose tissue, metabolism

## Abstract

The naked mole-rat (NMR) is a subterranean rodent known for its unique thermal biology, exceptional longevity and resistance to cancer and hypoxia. However, its thermal biology remains controversial, with various reports describing NMRs as poikilotherms, heterotherms, mesotherms or partial homeotherms. Here, we investigated whether the thermogenic potential of NMR brown adipose tissue and its UCP1 differ from those in mice and whether the lack of thermal insulation causes extreme changes in NMR body temperature upon cold exposure. Through longitudinal molecular, thermal, metabolic, and behavioral measurements, we found that NMRs initiated non-shivering thermogenesis and elevated body temperature but could not sustain it due to excessive heat loss and limits to substrate availability. Our results suggest that NMRs represent a unique thermoregulatory category that doesn’t fit neatly into traditional classifications. *In vitro* and *in vivo* experiments showed that the NMR UCP1 is functional and can be activated and inhibited as expected for most other mammals. We further demonstrated that artificial insulation can partially restore thermoregulatory capabilities in NMRs. This study employs an advanced methodology to characterize the thermal biology of NMRs and helps resolve a long-standing controversy in the field.

**Significance Statement:** This study provides critical insights into the thermal biology of naked mole-rats (NMRs), resolving long-standing controversies regarding their thermoregulatory strategies. We highlight the unique adaptations and limitations of NMR physiology by demonstrating that NMRs possess functional non-shivering thermogenesis and UCP1 but fail to maintain homeothermy due to excessive heat loss. Our findings suggest that artificial insulation can partially restore their thermoregulatory capabilities, offering a new perspective on the evolutionary and ecological significance of fur loss in NMRs. This research advances our understanding of mammalian thermal biology and presents an updated model for NMRs, bridging gaps between previous conflicting reports.

## Introduction

Naked mole-rats (NMRs; *Heterocephalus glaber*) hold a unique position among mammals, in part due to their unique thermal biology, exceptional longevity and resistance to cancer (1–3). These hairless creatures inhabit warm, humid burrows in tropical Northeast Africa, living in large colonies with a single breeding female (4). Their subterranean niche maintains a relatively stable temperature range of 29-32 °C throughout the year (5–8).

The thermoregulatory status of NMRs has been a subject of debate (9, 10), with various classifications proposed in the literature. These classifications often rely on four key terms (11, 12): homeothermic, poikilothermic, mesothermic, and heterothermic. Homeothermic animals maintain a relatively constant internal body temperature regardless of environmental conditions. Poikilothermy implies a state where internal body temperature varies significantly with ambient temperature in animals without effective autonomic temperature regulation (11). Mesothermic animals maintain body temperatures between those of typical homeothermic and poikilothermic animals (13, 14). Heterothermic animals can switch between homeothermic and poikilothermic strategies depending on environmental conditions or physiological state (12). Some researchers have suggested that NMRs, adapted to thermally buffered conditions, may have abandoned active thermoregulation, becoming thermoconformers to their environment (5, 15, 16). Despite this proposition, observations in the NMR colonial environment reveal homeothermy via behavioral thermoregulation, such as huddling and selection of warmer or colder burrow sectors (8, 10, 17, 18). However, when isolated from their stable habitat, individual NMRs demonstrate a complete dependence of their body temperature (T_b_) on the ambient temperature (T_a_), exhibiting a wide range of T_b_ across varying T_a_ tested (12–37 °C) (5, 6, 15). The complexity of NMR thermal biology is further emphasized by their varied classifications in the literature: poorly thermoregulating homeotherms (5, 6), endothermic poikilotherms (10, 15, 19–22), partial homeotherms (9), mesotherms (23), or heterotherms (24–26), This diversity of classifications underscores the unique and complex nature of NMR thermal biology.

Although mammals utilize shivering in order to boost heat generation upon cold exposure (27), the central role of heat production in small eutherian mammals is driven by more efficient non-shivering thermogenesis, which is managed by the sympathetic nervous system via noradrenergic innervation of brown adipose tissue (BAT) (28–31). The principal mechanism driving the production of heat in the BAT involves uncoupling protein 1 (UCP1), which resides in the inner mitochondrial membrane and dissipates the mitochondrial proton gradient in the form of heat (32). When the analysis of the first NMR genome assembly revealed that the UCP1 protein, despite being highly conserved across mammals, harbors unique changes within the fatty acid and nucleotide-binding domains, we hypothesized that these changes may reduce UCP1 uncoupling efficiency or regulation and result in diminished rates of thermogenesis (33, 34). However, recent investigations indicated a clear thermogenic potential in NMRs in response to cold stimuli (24, 25), challenging our previous assumptions and prompting further inquiry into their thermogenic mechanisms. Despite the presence of BAT and its innervation (35) and evidence of thermogenesis, including the detection of UCP1 (24, 25), comparative analyses of NMR with other mammals, such as mice, are lacking. Furthermore, despite suggestions that the inability to maintain stable T_b_ might be due to rapid heat dissipation (19, 36), the direct contributions of factors such as fur insulation and dietary lipids to NMR thermoregulation remains unexplored.

Here, we present our work aimed to elucidate the differences in thermogenic potential of NMR and mouse BAT and UCP1, and to investigate the impact of exogenous dietary lipid supplementation and thermal insulation on NMR thermoregulation. By employing a comparative approach, including the development of cell lines for the analysis of NMR UCP1 mutations and the introduction of fleece bed shelters to mimic fur insulation, we aim to uncover novel insights into NMR thermogenesis. Our findings reveal that NMRs possess functional non-shivering thermogenesis capabilities but fail to maintain homeothermy due to excessive heat loss, which can be partially mitigated by artificial insulation. This study offers the most comprehensive characterization of NMR thermogenic response available to date across a broad range of ambient temperatures.

## Results

### NMRs have BAT reserves and a functional UCP1

Non-shivering thermogenesis in mammals is supported by the UCP1 activity in the BAT. To understand the mechanisms underlying the failure of NMRs to maintain stable body temperature, we first examined the uncoupling properties of NMR UCP1 in comparison with those of its mouse ortholog. HEK 293 cells stably expressing NMR or mouse UCP1 displayed similar protein levels (Fig. 1*A*) localized in mitochondria (Fig. 1*B*). Upon activation by TTNPB, a retinoic acid analog that can specifically activate UCP1 but is not metabolized by cells (37, 38), both NMR and mouse UCP1 exhibited comparable oxygen consumption rates (OCR) and dose-dependent responses to modulators of cellular respiration (Fig. 1*C*), indicating functional similarity.

**Figure 1.**
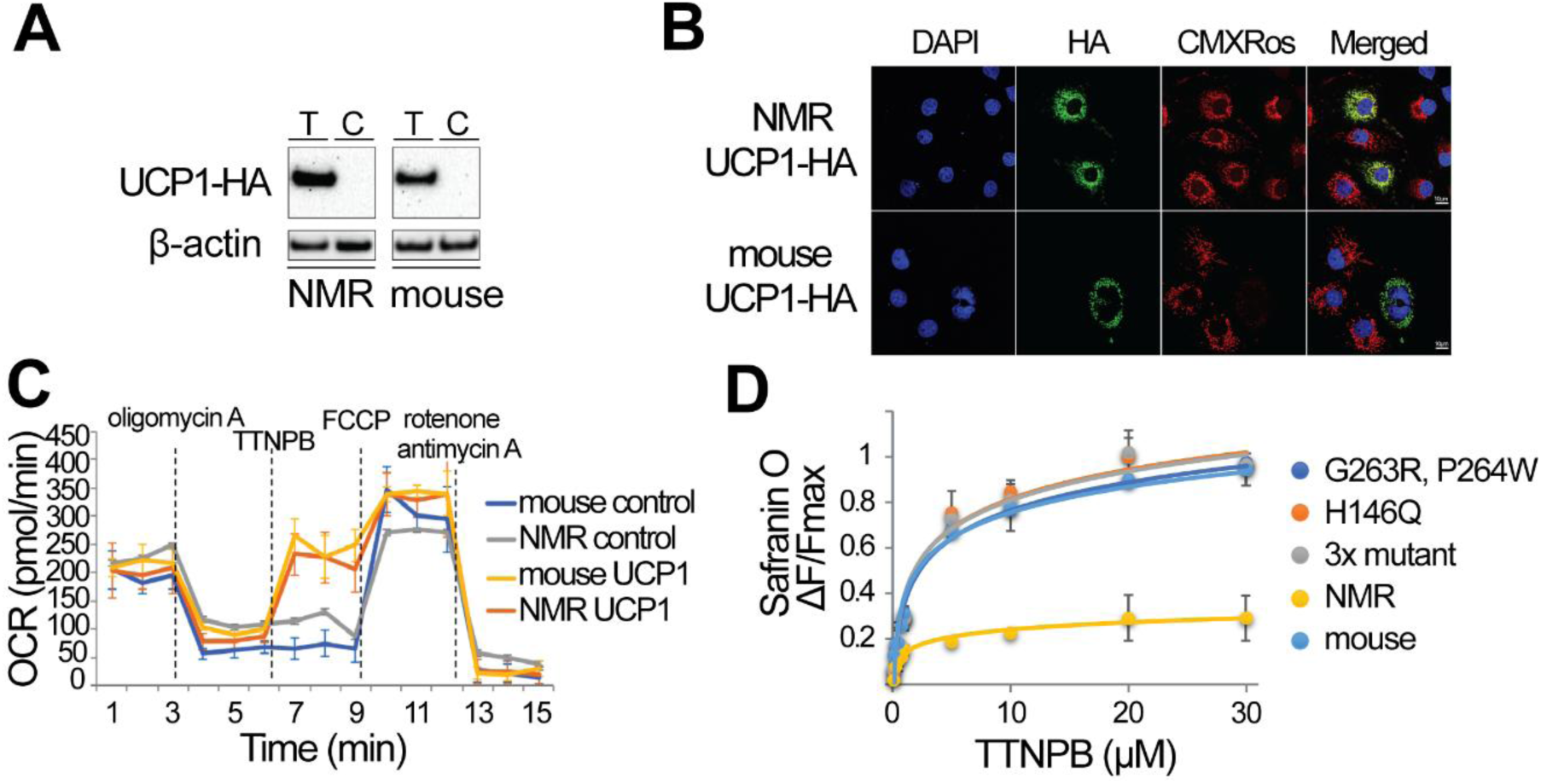
Characterization of NMR UCP1 function. (*A*) Western blotting of nmrUCP1-HA (left) and mUCP1-HA (right) proteins in stably transfected HEK 293 cells upon induction of expression with Tetracycline (+T) or Control (C). (*B*) Localization of NMR and mouse UCP1 to mitochondria. The proteins were immunostained based on their HA tag (green), mitochondria were stained with MitoTracker Red CMXRos (red), and nuclei with DAPI (blue). MitoTracker Red CMXRos is a cell-permeant probe that accumulates in active mitochondria and is retained after aldehyde fixation. The merged image shows colocalization of UCP1-HA and mitochondria, indicated by yellow areas. Note that not all cells show UCP1-HA signal due to variability inherent in the tetracycline-inducible expression system. Scale bar: 10 μm. (*C*) Oxygen consumption rates (OCR) measured by Seahorse assays in approximately 3×10^4^ plated HEK 293 cells expressing mouse or NMR UCP1-HA, demonstrating specific activation of proteins in comparison with control cells. The figure highlights basal respiration, proton leak respiration (after oligomycin treatment), specific UCP1 respiration (TTNPB), maximal substrate oxidation (FCCP) and non-mitochondrial respiration (antimycin A/rotenone). All values are mean ± SD from three independent experiments. (*D*) UCP1 activation in isolated HEK293 mitochondria (50 μg of mitochondrial protein). The graph shows changes in mitochondrial membrane potential in response to increasing concentrations of the UCP1 activator TTNPB. The y-axis (ΔF/Fmax) represents the change in safranin O fluorescence (ΔF) relative to the maximum fluorescence change (Fmax) observed upon complete mitochondrial depolarization with FCCP. Higher ΔF/Fmax values indicate greater depolarization, reflecting increased UCP1 activity. Mitochondria were isolated from HEK293 cells expressing NMR UCP1-HA, mouse UCP1-HA, or mouse UCP1-HA harboring NMR-specific mutations (G263R+P264W; H146Q; G263R+P264W+H146Q). All values are mean ± SD from three independent experiments.

NMR UCP1 differs from its mammalian orthologs by three amino acid residues at the conserved regulatory site, including a histidine important for proton transport (39) (Fig. S1). To assess their impact on UCP1 activity, we generated three mutant cell lines expressing mouse UCP1 with the following NMR-specific mutations: (1) G263R+P264W, (2) H146Q, and (3) G263R+P264W+H146Q. These specific mutations were chosen as they represent the key differences between mouse and NMR UCP1 sequences in highly conserved regions, potentially affecting protein function (33). Isolated mitochondria from these cells showed that natural NMR mutations did not diminish UCP1 activity compared to the wild-type mouse ortholog (Fig. 1*D*) and did not prevent NMR mitochondrial proton leak in response to TTNPB activation (Fig. 1*D*, Fig. S2*A*). While both NMR and mouse UCP1 showed activation in response to TTNPB, NMR UCP1 demonstrated a lower maximal activation capacity compared to wild-type and mutant mouse UCP1. To verify this, we conducted additional safranin O quenching experiments using Palmitate as a UCP1 activator. These experiments revealed a trend similar to TTNPB, with Palmitate inducing UCP1 uncoupling (Fig. S2C). Both TTNPB and Palmitate experiments suggest that NMR UCP1-HA in HEK293 mitochondria had a lower maximal activation capacity compared to mouse UCP1, including variants with NMR-specific mutations. This consistent pattern across different activators suggests functional differences between NMR and mouse UCP1 when expressed in vitro. Interestingly, while whole-cell OCR showed similar responses for mouse and NMR UCP1 (Fig. 1C), isolated mitochondria membrane potential measurements revealed lower activity for NMR UCP1 compared to mouse UCP1 (Fig. 1D). This apparent discrepancy between whole-cell and isolated mitochondria experiments highlights the complexity of comparing UCP1 function across different experimental systems and is addressed in detail in the Discussion.

We then investigated the effect of GDP, a known inhibitor of UCP1 (40). GDP caused a dose-dependent inhibition of both mouse and NMR UCP1 activity (Fig. S2*A*). All UCP1 variants showed similar inhibition profiles (Fig. S2*B*), suggesting that these three sequence changes do not affect the GDP-binding site and inhibitory mechanism of NMR UCP1.

To validate these findings in a more physiologically relevant context, we examined the potential for UCP1-mediated non-shivering thermogenesis by analyzing the BAT. Post-mortem anatomical inspection confirmed the presence of BAT depots in NMRs, with anatomical locations similar to those in mice (Fig. 2*A*). Histological analyses of BAT sections also showed similar morphological characteristics of brown adipocytes including multilocular lipid droplets (Fig. 2*B*). UCP1 expression (Fig. S2*D*) supported the thermogenic potential of NMR BAT (24, 25).

**Figure 2.**
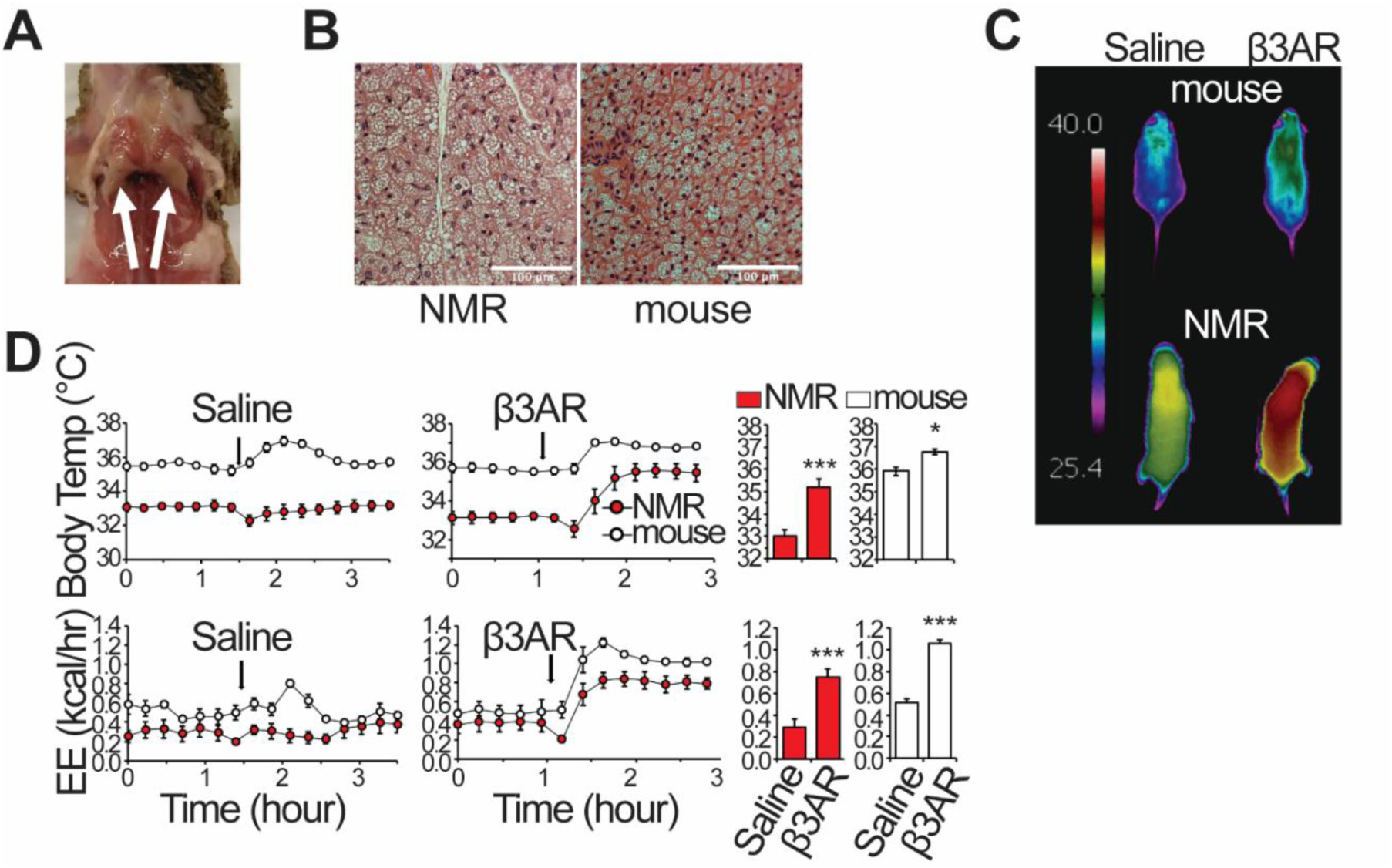
Endogenous thermogenesis in the NMR. (*A*) Imaging of NMR showing two interscapular brown fat depots. (*B*) Histology of NMR and mouse interscapular brown adipose tissue, staining with H&E. Scale bar, 100 μm. (*C*) Thermal images of NMR and mouse surface temperature in response to saline (control) and β3-adrenergic agonist CL-316,243 interventions. (*D*) Indirect calorimetry monitoring of NMRs and mice in response to saline and β3-adrenergic agonist CL-316,243 interventions (β3AR). Upper panel: NMR and mouse average core body temperature. Lower panel: NMR and mouse average energy expenditure. Data points were collected every 12 minutes. Column graphs represent means of the entire data collection period following each injection, starting from the first post-injection data point until the end of the monitoring period. All values are mean ± SEM from five mice and four NMRs (N = 5 mice, N = 4 NMRs). All bar plots reach statistical significance (* p <0.05, *** p< 0.001).

We then examined UCP1 function in mitochondria isolated directly from BAT of mice and NMRs. Consistent with our data on mitochondrial membrane potential (41) measurements in HEK293 cells, both mouse and NMR BAT mitochondria showed UCP1 activation in response to TTNPB and inhibition by GDP (Fig. S2*E*). Interestingly, the magnitude of activation was notably higher in BAT-derived mitochondria compared to HEK293-expressed UCP1, particularly for NMR UCP1. This apparent difference is addressed in detail in the Discussion section, highlighting the complexity of comparing UCP1 function across different experimental systems. The similar TTNPB-activated, GDP-inhibited uncoupling activity in both species is consistent with functional non-shivering thermogenesis capability in NMRs. Taken together, these results demonstrate that, like in mice, the BAT in NMRs contains functional UCP1 and exhibits a clear thermogenic potential.

### Active thermogenic response following pharmacological stimulation of brown fat in NMRs

These properties of BAT and UCP1 led us to reevaluate the ability of NMRs to regulate body temperature *in vivo.* In response to cold exposure, the mammalian sympathetic nervous system releases noradrenaline to stimulate the β-adrenergic receptors (β3AR) (28, 32). These receptors are primarily localized within adipose tissues, and their activation coordinates the release of fatty acids from white adipose tissue for thermogenesis in BAT (42). When injected with a specific β3AR agonist CL-316,243, non-anesthetized freely moving NMRs housed at thermoneutral conditions demonstrated a profound 2.5 °C increase in surface temperature determined non-invasively by thermal imaging (Fig. 2*C*). The corresponding increase in mice appeared lower, around 1 °C (Fig. 2*C*), yet the changes in both species were significant (p<0.05) compared to saline administration (Fig. S3*A*).

To more precisely characterize BAT thermogenic activity, we monitored metabolic rate and core body temperature in mice and NMRs implanted with wireless thermal sensors. To separate the non-specific effects of stress response caused by injection, we initially monitored all animals during an intraperitoneal injection of saline. We observed a transient increase in body temperature and metabolic rate in mice, but no change in respiratory exchange ratio (RER) (Fig. 2*D* and S3*B*). In contrast, NMRs showed no apparent physiological response to saline injection (Fig. 2*D*). Upon β3-adrenergic stimulation (1 mg/kg CL-316,243 (CL)), both mice and NMRs exhibited a robust thermogenic response characterized by increased body temperature, increased metabolic rate, and increased lipid oxidation as indicated by a lower RER (Fig. 2*D* and S3*B*). Strikingly, NMRs showed a greater absolute and fold increase in core body temperature compared to mice. Overall, NMRs responded to an acute thermo-mimetic stimulus in a manner qualitatively similar to that of mice, supporting their capacity for active thermogenesis during cold exposure.

### Exogenous substrate helps maintain NMR body temperature during cold exposure

The capacity for UCP1-mediated non-shivering thermogenesis in NMRs seems at odds with their reported thermally labile response to cold exposure. To address variability in older studies reporting NMR body temperature across a broad range of ambient temperatures (5, 6, 15), we performed continuous longitudinal measurements of NMR core T_b_ over an ambient temperature range of 18-37 °C. We confirmed that NMRs fail to maintain euthermia (Fig. 3*A*).

**Figure 3.**
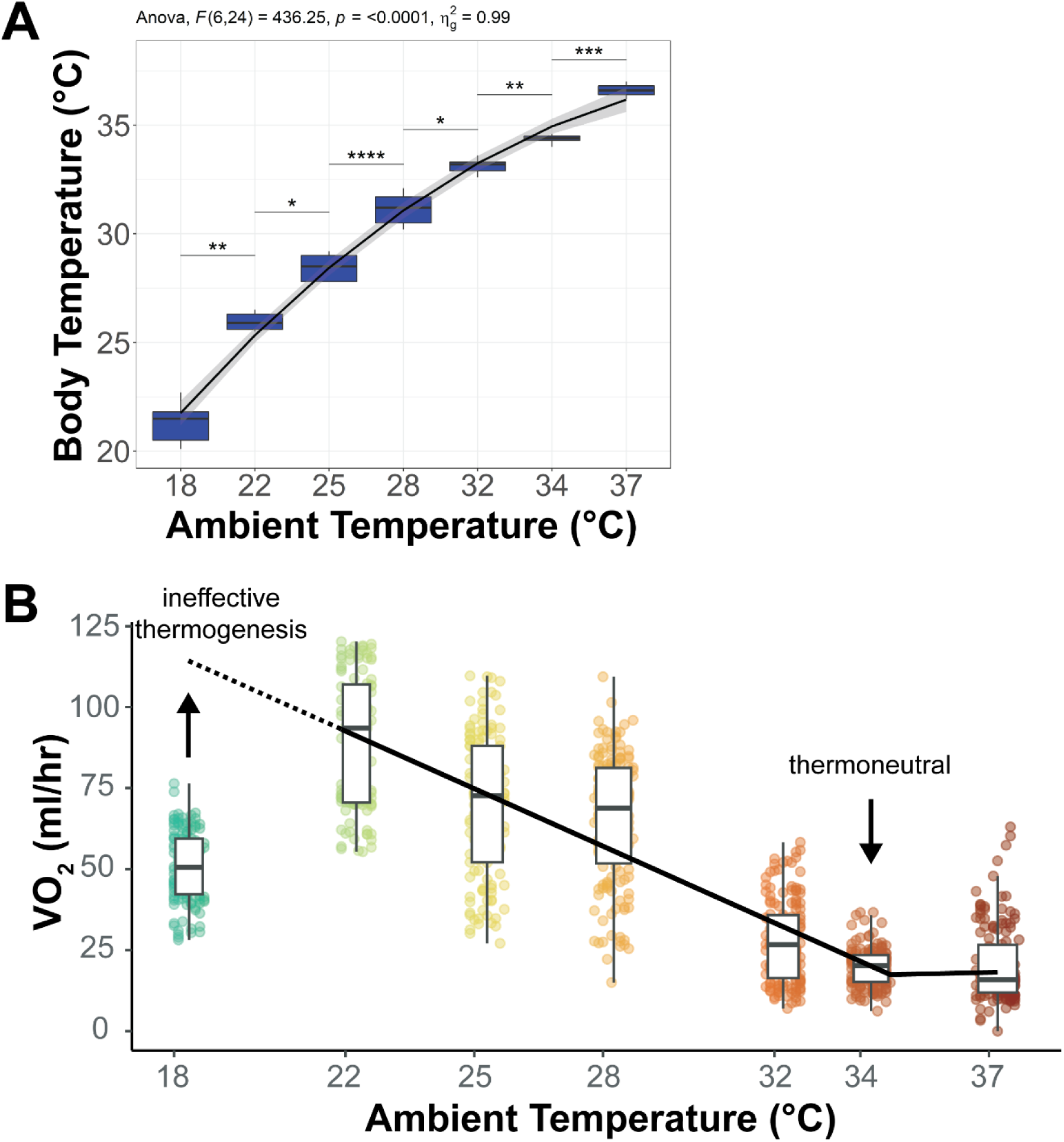
Relationship between ambient temperature, core body temperature, and metabolic rate in naked mole-rats (NMRs). See also Table S1. (*A*) Core body temperature of individual non-anesthetized NMRs as a function of ambient temperature. Each box represents the distribution of data from five individual NMRs (N=5) at each ambient temperature. Core body temperatures were measured continuously using implanted temperature sensors as ambient temperature was varied from 18°C to 37°C. (*B*) Scholander plot of metabolic rate vs ambient temperature in free-moving, non-anesthetized, non-insulated NMRs. Each point represents the mean metabolic rate at each 12 minute interval at the indicated ambient temperature (N=5 NMRs per temperature). The solid line shows the linear relationship between metabolic rate and ambient temperature from 22°C to 34°C. The body temperature of 34°C was revealed as thermoneutral temperature, where metabolic rate is lowest and begins to plateau. Below 22°C, NMRs show signs of ineffective thermogenesis, as indicated by the deviation from linearity.

To accurately determine the thermoneutral temperature for NMRs, we employed a Scholander plot analysis (Fig. 3*B*). In this plot, metabolic rate is graphed against ambient temperature. The thermoneutral zone is identified as the range of ambient temperatures where the metabolic rate is at its lowest and relatively constant. Below this zone, metabolic rate increases linearly with decreasing temperature as the animal expends more energy to maintain body temperature. Above this zone, metabolic rate may increase due to the energetic costs of dissipating heat. Our analysis revealed that the thermoneutral temperature for individually housed NMRs is 34°C, higher than that previously reported for animals housed in groups (43).

Given the functional UCP1 and BAT reserves demonstrated both here and recently (24, 25), and the potential to generate heat and elevate body temperature, NMRs’ inability to defend their T_b_ remains unclear. One plausible mechanism is a lack of sufficient fuel for thermogenesis. To test this possibility, we measured the body temperature of NMRs during cold exposure (22 °C) and tested the effect of exogenous lipid substrates. Dietary olive oil supplementation allowed NMRs to maintain body temperature 2.6±1.6 °C (95% CI) higher than controls at 22 °C (Fig. 4*A*). In contrast, C57BL/6J mice showed no effect on their body temperature upon 4 °C cold challenge with olive oil supplementation (Fig. 4*B*). While these results suggest that exogenous lipids help NMRs maintain a higher body temperature during cold exposure, it’s important to note that this observation alone does not definitively prove increased lipid oxidation in BAT. We also measured the concentration of plasma non-esterified fatty acids (NEFA) in NMRs before, during, and after the cold-challenge. In response to 22 °C cold exposure, NMR blood levels of free fatty acids increased twofold (Fig. 4*C*), indicating an effort to fuel BAT thermogenesis. Together, these data suggest that NMRs mobilize energy stores from white adipose tissue through lipolysis in response to cold, and exogenous substrates like olive oil help maintain a higher body temperature, likely by increasing fatty acid uptake into BAT.

**Figure 4.**
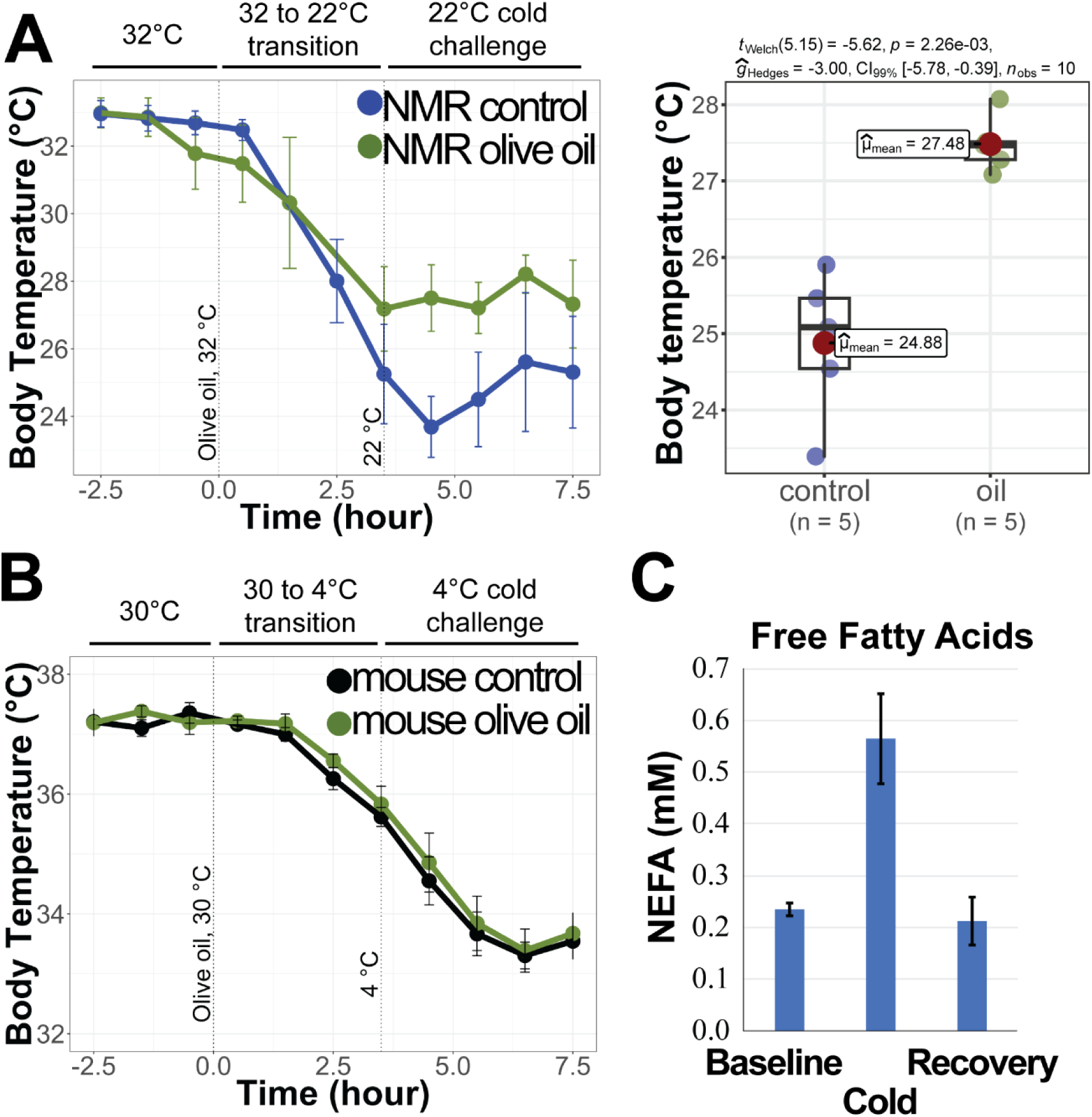
Effect of olive oil supplementation on body temperature during cold challenge. (*A*) NMR body temperature during cold challenge from Ta=32 to Ta=22 °C. Olive oil was administered by gavage to animals maintained at 32°C. The ambient temperature was then transitioned to 22°C over 3 hours at a constant rate. The boxplot represents the mean body temperature during the entire stable temperature period at 22°C, excluding the temperature transition phases. N = 5 NMRs. (*B*) Mouse body temperature during cold challenge from Ta=30°C to Ta=4°C. N = 5 mice. (*C*) Plasma free fatty acid levels in NMRs at baseline, upon exposure to 22 °C for 7 hours, and following recovery for 72 h (mean±SD, n=2). See also Table S2.

### Excessive heat loss challenges homeothermy in mice

Exogenous substrate supplementation, functional lipolysis and robust non-shivering thermogenesis help maintain NMR body temperature during cold exposure, but these factors do not explain the absence of homeothermy. NMR skin, known for its loosely folded morphology, well-developed vascularization of the upper dermis, high thermal conductivity, and poor insulation properties due to the absence of pelage (36, 44), likely contributes to body temperature drops during cold challenges by facilitating heat loss through the hairless skin (24, 25, 36).

We first examined the role of insulation by comparing metabolic rate and core body temperature in shaved versus unshaved (furry) C57BL/6J mice exposed to ambient temperatures below thermoneutrality (6-28 °C). This revealed the remarkable metabolic plasticity and cold adaptation in this species (28, 45). Even at the coldest ambient temperature of 6 °C, furry mice only experienced a 1.1±0.4 °C (95% CI) drop in core body temperature (Fig. 5*A, B*). They achieved euthermia by doubling energy expenditure (EE) (Fig. 5*C, D*) and increasing fatty acid oxidation, as reflected by lower RER values (Fig. 5*E, F*). In contrast, shaved mice experienced a significant (2.1±0.3 °C, 95% CI) drop in core body temperature and a threefold increase in metabolic rate, highlighting the role of fur insulation. Although the respiratory exchange ratio of shaved mice was higher (indicating less fatty acid oxidation) at thermoneutrality, it remained unchanged during the cold challenge. During the recovery days at thermoneutrality, shaved mice exhibited RER values greater than 1.0, indicating *de novo* lipogenesis and rapid energy conversion to replenish adipose tissue stores (Fig. 5*E, F*).

**Figure 5.**
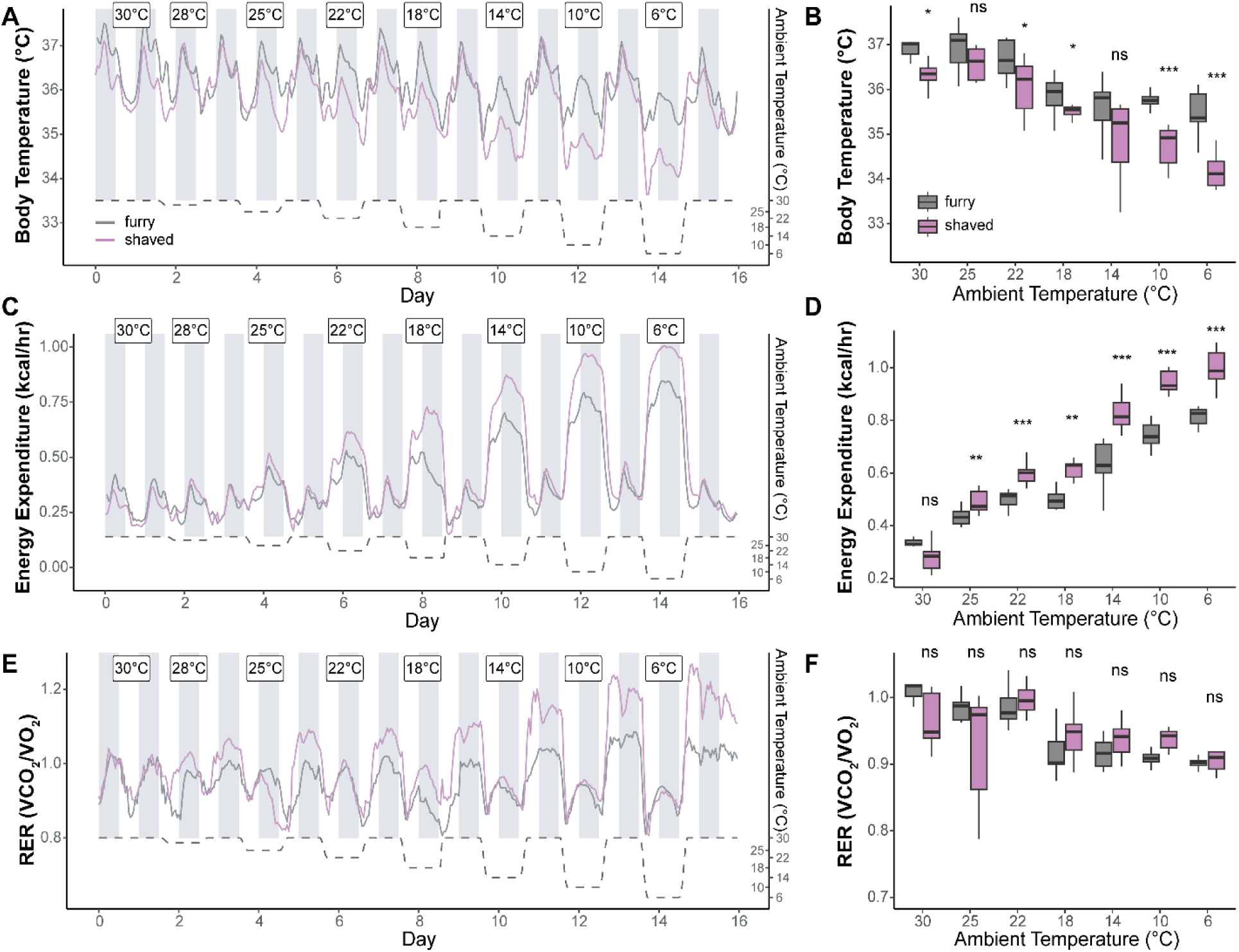
Impact of fur insulation on thermoregulation and metabolism in mice exposed to various ambient temperatures. (*A*) Continuous core body temperature measurements of furry (gray) and shaved (pink) mice during exposure to decreasing ambient temperatures from 30°C to 6°C. Each temperature condition lasted 24 hours, alternating with 24-hour recovery periods at thermoneutrality (30°C). Ambient temperature is indicated by the dashed line (right y-axis). (*B*) Box plot comparison of core body temperatures in furry and shaved mice at each ambient temperature. Data represent dark phase measurements. (*C*) Continuous energy expenditure measurements of furry and shaved mice during the temperature challenge protocol described in (A). (*D*) Box plot comparison of energy expenditure in furry and shaved mice at each ambient temperature. Data represent dark phase measurements. (*E*) Continuous respiratory exchange ratio (RER) measurements of furry and shaved mice during the temperature challenge protocol. (*F*) Box plot comparison of RER in furry and shaved mice at each ambient temperature. Data represent dark phase measurements. For all panels, N = 9 mice per group. In box plots, the central line indicates the median, box limits represent upper and lower quartiles, and whiskers extend to 1.5 times the interquartile range. Statistical significance between furry and shaved groups at each temperature is indicated as follows: ns - not significant, * p < 0.05, ** p < 0.01, *** p < 0.001. See Table S3 for detailed statistical analysis.

### Effect of insulation on NMR thermoregulation and metabolism

We next investigated how NMRs regulate body temperature in response to ambient temperatures below thermoneutrality. Continuous monitoring of NMR core body temperature, metabolic rate, and substrate utilization in response to cold ambient temperatures revealed an endothermic response. Each transition from thermoneutrality to a colder ambient temperature (18-28 °C) triggered a robust increase in metabolic rate and increased fatty acid oxidation (decreased RER values). Despite these adaptations, core body temperature closely mirrored ambient temperature changes (Fig. 6*A*-*F*).

**Figure 6.**
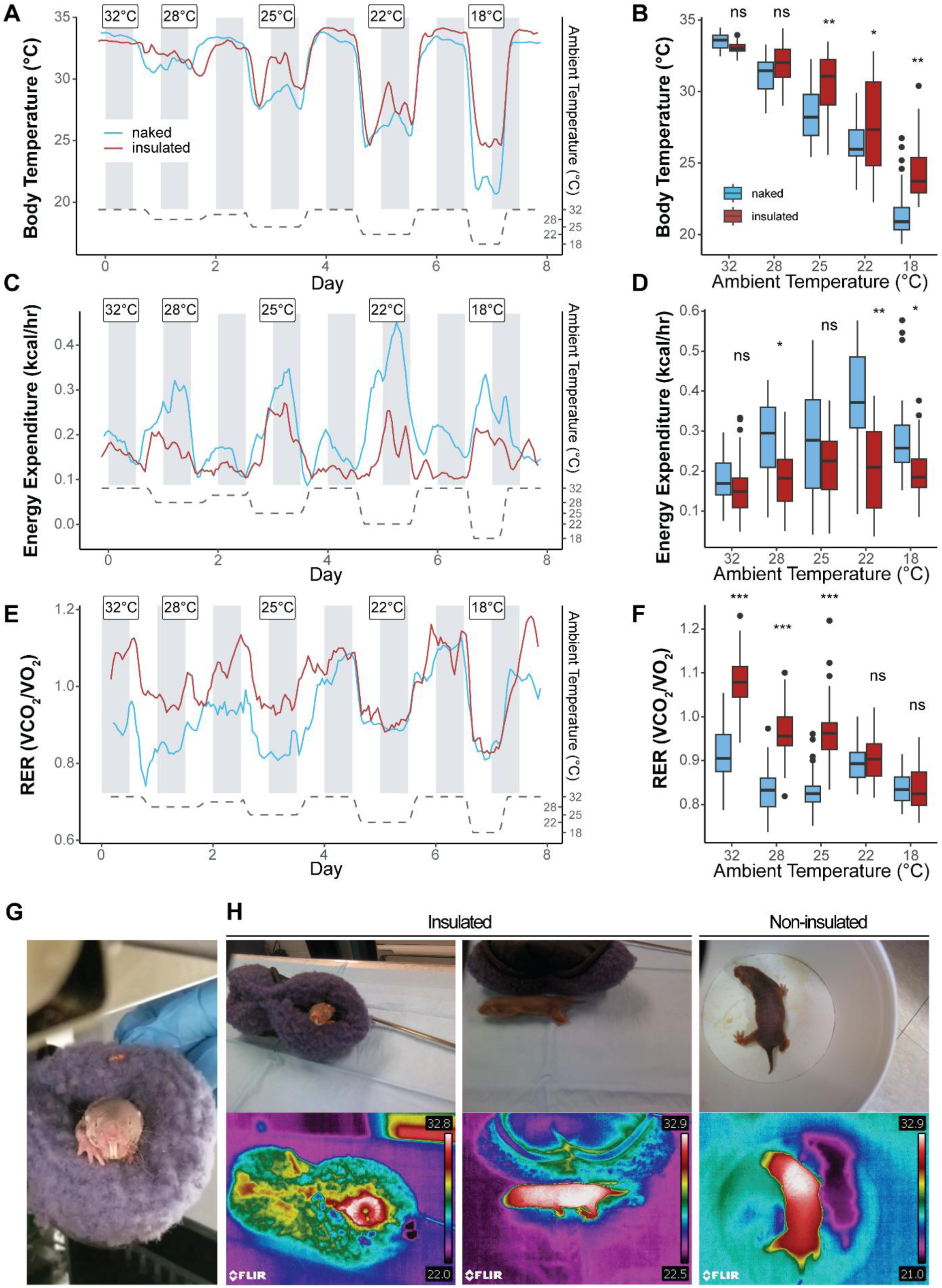
Impact of artificial insulation on thermoregulation and metabolism in naked mole-rats (NMRs) exposed to various ambient temperatures. (*A*) Continuous core body temperature measurements of non-insulated (blue) and insulated (red) NMRs during exposure to decreasing ambient temperatures from 32°C to 18°C. Each temperature condition lasted 24 hours (12 hours for 18°C), alternating with 24-hour recovery periods at 32°C. Ambient temperature is indicated by the dashed line (right y-axis). (*B*) Box plot comparison of core body temperatures in non-insulated and insulated NMRs at each ambient temperature. (*C*) Continuous energy expenditure measurements of non-insulated and insulated NMRs during the temperature challenge protocol described in (A). (*D*) Box plot comparison of energy expenditure in non-insulated and insulated NMRs at each ambient temperature. (*E*) Continuous respiratory exchange ratio (RER) measurements of non-insulated and insulated NMRs during the temperature challenge protocol. (*F*) Box plot comparison of RER in non-insulated and insulated NMRs at each ambient temperature. For panels A-F, data represent the entire stable temperature period after the target temperature was reached. Temperature transition periods (3 hours) between different ambient temperatures were excluded from the analysis. N = 5 NMRs per group. In box plots, the central line indicates the median, box limits represent upper and lower quartiles, whiskers extend to 1.5 times the interquartile range, and individual points represent outliers. Statistical significance between non-insulated and insulated groups at each temperature is indicated as follows: ns - not significant, * p < 0.05, ** p < 0.01, *** p < 0.001. See Table S4 for detailed statistical analysis. Hourly average body temperature values at specified ambient temperatures can be found in Supplementary Figure S5. (*G*) Photograph of an NMR using a fleece shelter as artificial insulation. (*H*) Thermal images of NMRs’ surface temperature in response to room temperature (22°C) challenge. Left panel: NMR in fleece shelter. Middle panel: NMR immediately after leaving an insulated enclosure. Right panel: Non-insulated NMR after 2 hours at room temperature. Color scale indicates surface temperature in °C.

To gain insights into the effect of insulation on NMR thermoregulation and metabolism, we introduced fleece bed shelters to help retain generated body heat (Fig. 6*G*, Fig. S*4*). When housed at 28 °C, body temperature dropped similarly in insulated (by 2±0.4 °C) and non-insulated (by 2±0.6 °C, 95% CI) NMRs (Fig. 6*A, B*). However, insulation reduced the metabolic demand for heat production. Insulated NMRs increased EE by 36%, while non-insulated animals increased EE by 83% (Fig. 6*C, D*). Similarly, insulated NMRs utilized less fatty acid oxidation than non-insulated animals (Fig. 6*E, F*). This trend continued at 25 °C, with a further increase in metabolic rate in both groups. Insulated NMRs maintained higher body temperatures and lower metabolic rates than non-insulated animals, suggesting a lower metabolic demand. Insulated NMRs could maintain euthermia at 25 °C by inducing thermogenesis, fatty acid oxidation, and elevating of EE.

When exposed to 18 and 25 °C ambient temperatures, insulated NMRs had significantly higher body temperatures than non-insulated animals (Fig. 6*A, B*). Moreover, their observed maximum T_b_ values were within the range detected for NMRs housed at thermoneutrality (Fig. S5), indicating the capacity for homeothermy.

Thermal imaging data of insulated NMRs exposed to an ambient temperature of 22 °C showed that they retained generated heat more effectively, maintaining their surface temperature at 32.6±0.6 °C (Fig. 6*H*). Corresponding measurements of non-insulated NMRs confirmed the endogenous non-shivering thermogenesis in BAT, indicated by the interscapular surface temperature of 8.7±2 °C higher than ambient temperatures. Thermal images from cold-induced thermogenesis also revealed heat loss through the skin, resulting in much lower temperatures of the rest of the body surface (Fig. 6*H*, right panel). Specifically, the surface temperature of extremities appeared almost as low as the ambient temperature. These findings indicate that while NMRs possess functional BAT capable of non-shivering thermogenesis, excessive heat loss through their skin precludes homeothermy, which can be partially mitigated by artificial insulation.

## Discussion

Our study aimed to investigate the ability of naked mole-rats (NMRs) to regulate body temperature through non-shivering thermogenesis in response to cold exposure and to determine the factors that preclude homeothermy. Our molecular, thermal, metabolic and behavioral findings demonstrated that NMRs could initiate robust non-shivering thermogenesis and elevate body temperature when exposed to cold. However, they failed to sustain this elevated temperature due to excessive heat loss through their skin.

Notably, providing artificial insulation partially restored their ability to maintain homeothermy. This significant improvement in heat retention indicates that the primary challenge for NMR thermoregulation is not a lack of thermogenic capacity but rather their inability to prevent heat loss. This finding challenges previous assumptions about NMR thermoregulation and highlights the importance of considering both heat generation and retention in understanding their thermal biology.

One of key findings of our study is the revision of the thermoneutral temperature for naked mole-rats (NMRs). Our Scholander plot analysis revealed that the true thermoneutral temperature for individually housed NMRs is 34°C, higher than previously reported for group-housed animals (43). This finding has important implications for understanding NMR thermoregulation, suggesting that individual NMRs may experience greater thermoregulatory stress at temperatures previously considered thermoneutral. The linear relationship between metabolic rate and ambient temperature from 22°C to 34°C indicates that within this range, NMRs can effectively adjust their metabolism to cope with temperature changes. However, at 18°C, we observed ineffective thermogenesis, indicating a lower limit to their thermoregulatory capacity. This revised understanding necessitates a reevaluation of previous studies on NMR thermal biology, as experiments conducted at 32°C may represent mild cold stress for individually housed NMRs.

Previous studies suggested that the lack of insulating fur as a factor contributing to NMR inability to maintain body temperature during cold exposure (6, 19, 20, 24, 36, 44), but they did not confirm this experimentally. Withers and Jarvis’s study even appeared to disprove this by showing no difference in body temperature between artificially insulated (in the form of nesting material) and non-insulated NMRs at 25 and 30 °C, despite a decreased metabolic rate in the former ones (5). Our results differed, likely due to the more effective insulation provided by the shelters in our experiments, which covered a larger body surface area and significantly reduced heat loss.

Our findings provide new insights into the ongoing debate about NMR thermoregulatory classification. The initial attempts to characterize NMR metabolic response to cold, conducted by McNab in 1966 (6) and Withers and Jarvis in 1980 (5), described NMR as a poorly thermoregulating homeotherm. However, Buffenstein and Yahav’s 1991 study (15) challenged this conception revealing a predominantly poikilothermic behavior of NMR below the thermoneutral zone. Two of these three studies (5, 15) showed that the metabolic response of the NMR to cold differed from a typical mammalian pattern. As fairly suggested in the recent review (19), where the NMR was also being referred to as a poikilotherm, it was likely that the amount of measurements conducted in the McNab’s study was not sufficient to reveal the details of NMR energetics. The most recent studies proposed that NMRs should rather be considered partially homeothermic (9), mesothermic (23) or heterothermic (24–26), but not poikilothermic. Different classifications proposed by these studies illustrate the magnitude of this controversy.

While our findings support previous observations of NMR metabolic responses to cold (15)), which characterized them as endothermic poikilotherms, we question the basis for NMR poikilothermy. The definition of poikilothermy implies a large variability of body temperature as a function of ambient temperature in organisms without effective autonomic temperature regulation (11). Despite showing variability in body temperature with environmental changes, NMRs exhibited robust thermogenic capabilities and body temperature elevation. Moreover, poikilotherms are usually known for their ability to survive low or even freezing body temperatures, whereas NMRs develop pathological phenotypes such as necrosis of the extremities when their body temperature drops to 20 °C and lower for prolonged periods of time (20, 21).

In support of (6, 23, 24) but in contrast to (15), our study demonstrated a homeothermic response in which NMR increased oxygen consumption as ambient temperature decreased from 32 to 22 °C. While typical poikilotherms show a decrease in metabolic rate that allows them to survive through cold, NMRs experience pathological challenges when their body temperature drops below 20 °C, including the challenge of performing metabolic biochemical reactions with a core temperature of 21 °C. This suggests that, in response to cold exposure, individually housed NMRs exhibit characteristic features of homeotherms that fail to defend their body temperature, rather than that of true poikilotherms.

Given NMRs ability to metabolically produce heat together with inability to defend their body temperature could be seen as characteristics that align them with mesotherms. Although this term is less commonly used and somewhat controversial in the field, mesotherms is thought to maintain their body temperatures higher than the environment but lower and more variable than true homeotherms decreasing their metabolic rates as ambient temperature drops (13, 14). However, NMR thermoregulatory strategy seems more complex than simply maintaining a metabolic rate intermediate between that of endotherms and ectotherms. Moreover, the term “mesothermic” is typically applied to large aquatic animals without specialized thermogenic tissues, not small terrestrial mammals.

While NMRs do show some ability to regulate their body temperature under certain conditions, it’s not clear if they exhibit the distinct switches between homeothermy and poikilothermy in response to environmental conditions or physiological state – characteristic feature of true heterotherms (12). Their thermoregulatory switches are not as distinct or regular as those seen in classic heterotherms (e.g., daily torpor or seasonal hibernation). NMR thermoregulatory strategy is more of a constant blend rather than a clear switch between distinct states. Based on our findings, we propose that NMRs represent a unique thermoregulatory category that doesn’t fit neatly into traditional classifications.

We previously hypothesized that specific amino acid changes in NMR UCP1 might contribute to their unique response to cold. Multiple species, including pigs, horses, elephants, and whales, harbor large deletions of the UCP1 locus, resulting in inactive or poorly developed brown adipose tissue (BAT) and defective thermal regulation in young animals (46, 47). To investigate whether NMR UCP1 was a causal factor in the observed cold sensitivity, we reevaluated the functional state of NMR UCP1 *in vitro* and *in vivo,* as well as the response of BAT to cold exposure and β3AR activation. Our molecular and cellular studies revealed that NMR UCP1 is functional and can be both activated and inhibited, similar to UCP1 from other mammals. However, we observed an interesting discrepancy between whole-cell oxygen consumption rates (Fig. 1C) and isolated mitochondria membrane potential measurements (Fig. 1D) when comparing mouse and NMR UCP1 function. While whole-cell OCR showed similar responses, isolated mitochondria experiments indicated lower activity for NMR UCP1. This apparent contradiction can be explained by several factors. First, safranin O quenching assay used for isolated mitochondria directly measures membrane potential and is likely more sensitive to subtle differences in uncoupling activity compared to whole-cell OCR measurements. Second, in whole cells, factors such as substrate availability and delivery to mitochondria may be rate-limiting, potentially masking differences in UCP1 activity that become apparent in isolated mitochondria where these factors are controlled. Third, OCR experiments were normalized to cell number, while safranin O quenching was normalized to mitochondrial protein amount. This could amplify differences if there are variations in mitochondrial content or UCP1 expression levels between samples. Our study also revealed interesting differences in UCP1 function between heterologous expression in HEK293 cells and native expression in BAT mitochondria. While both systems demonstrated functional UCP1 activity, the magnitude of activation, particularly for NMR UCP1, was notably higher in BAT-derived mitochondria. While previous studies have shown that HA-tagged UCP1 generally retains its activity (49, 50), we cannot rule out the possibility that the tag affects UCP1 function, particularly in a species-specific manner. These observations highlight the complexity of performing comparative analyses of mouse and NMR UCP1 activity outside their natural physiological context. While heterologous expression systems provide valuable insights, they may not fully recapitulate the complexity of the native environment. It’s worth noting that primary NMR cell cultures do not grow well at 37°C and are normally maintained at 32°C, which is closer to their natural environment. This temperature difference could also contribute to the observed discrepancies and underscores the challenges in directly comparing UCP1 function between species with different physiological adaptations.

Thus, unlike swine, NMR UCP1 is functional and could be both activated and inhibited, as expected for most other mammals. NMR BAT morphology had pronounced multilocular lipid droplets, typical of rodent BAT. The NMR used in this study had a body composition and body mass similar to that of mice as shown by nuclear magnetic resonance (Fig. S6). Curiously, unlike the striking adaptation of mice to develop additional thermogenic capacity in response to short-term cold or adrenergic activation, NMRs did not show improved thermal responses even after one year of moderate cold exposure (20), highlighting distinct differences from other rodents that can adapt their thermogenic capacity. The ability of NMRs to upregulate body temperature and metabolic rate *in vivo* by pharmacologically stimulating β3AR confirmed that NMRs have intact thermogenic physiology (24), as in other rodents (28). This finding in conscious animals is consistent with the results of the NMR response to noradrenaline under anesthesia (16).

While our study provides comprehensive insights into NMR thermoregulation, it also has limitations. The artificial insulation used in our experiments may not perfectly insulate NMRs throughout cold exposure as animals were free enter and leave the insulated area voluntarily. While this approach minimized stress on the animals and better reflected their natural burrowing behavior, it could also lead to decreased average body temperatures and potential underestimation of the full effect of insulation. Future studies could address this by developing methods to provide more consistent insulation while minimizing stress on the animals. Despite these limitations, the significant differences observed between insulated and non-insulated conditions strongly suggest that the fleece shelters provided substantial thermoregulatory benefits. Another limitation of our study is the lack of direct measurement of UCP1 expression changes in response to cold exposure and insulation. While our experimental design aimed to mitigate potential cold-induced expression changes by returning animals to thermoneutrality between cold challenges, we cannot rule out the possibility that UCP1 expression was altered during the course of our experiments. Changes in UCP1 expression could potentially affect the interpretation of our results, as increased UCP1 levels could contribute to improved cold tolerance independent of the insulation effects we observed. Future studies should directly measure UCP1 expression changes in NMRs under various thermal conditions and insulation scenarios. Such studies could help distinguish between the effects of insulation and potential adaptive changes in UCP1 expression, further elucidating the mechanisms underlying NMR thermal biology. Another limitation of our study was the inability to measure blood glucose levels in cold-exposed NMRs. Recent research has shown that mouse BAT takes up significant amounts of glucose upon cold exposure, suggesting an important role for glucose metabolism in thermogenesis. However, the challenges of blood collection in NMRs without causing stress or disrupting continuous metabolic measurements prevented us from exploring this aspect of NMR thermal biology. For the same reason, we chose not to directly measure circulating NEFA levels after lipid gavage and during cold exposure. While we assumed an increase in circulating fatty acids based on previous studies in other rodents, direct measurements would have provided stronger evidence for our hypothesis about NEFA as a limiting factor during cold exposure in NMRs. Future studies could address this limitation by developing minimally invasive techniques for blood glucose and circulating lipids monitoring in NMRs. To further elucidate the mechanism of lipid-induced thermogenesis in NMRs, future studies should investigate whether the thermogenic effect of lipid gavage can be prevented by beta-receptor antagonists. This approach would provide important mechanistic insights into the role of beta-adrenergic signaling in NMR thermogenesis and its interaction with lipid availability. Lastly, our study has focused primarily on UCP1-mediated non-shivering thermogenesis in naked mole-rats. However, the complexity of mammalian thermoregulation suggests that other thermogenic mechanisms may play important roles in these unique animals. Future studies should investigate alternative thermogenic pathways, including those mediated by the adenine nucleotide translocator, basal proton leak, and shivering thermogenesis.

Our results highlight the importance of considering NMR thermal biology in the broader context of mammalian physiology and evolution. An intriguing implication of our study is the potential evolutionary advantage of fur loss in NMRs. Living in warm, densely populated burrows, NMRs may benefit from enhanced heat transfer through skin-to-skin contact, reducing individual thermal demands (43). This adaptation may parallel the evolutionary trajectory of humans. A long-standing hypothesis for the loss of body hair in humans is that once humans were able to maintain euthermia with clothing, shelter, and fire, the detrimental characteristic of fur as a welcoming refuge for disease-carrying parasites made loss of body hair advantageous (48). Since both humans and NMRs achieve euthermia in their respective environments, nakedness in these species appears to be a case of convergent evolution.

In conclusion, we have demonstrated that NMRs possess robust non-shivering thermogenesis and the ability to elevate body temperature, although excessive heat loss precludes sustained homeothermy. Providing artificial insulation can partially restore their thermoregulatory capabilities. Our study advances the understanding and presents an updated model of NMR thermal biology with the hope of resolving the long-standing controversy in the field. The thermal biology of NMRs represents a fascinating example of how evolutionary adaptations to a specific niche can lead to unique physiological traits, challenging our traditional categorizations of thermoregulatory strategies in mammals.

## Materials and Methods

### Animal ethics

Animal experiments were carried out according to the protocols approved by the Institutional Animal Care and Use Committee (IACUC) of the Brigham and Women’s Hospital, Harvard University, University of Illinois-Chicago and University of Rochester.

### Generation of UCP1-expressing HEK293 cell lines

For the analysis of NMR and mouse UCP1 in HEK 293 cells, mouse UCP1 CDS (GenBank: BC012701.1) and NMR UCP1 mRNA isolated from frozen tissues were transcribed to cDNA and cloned into the pcDNA™5/FRT/TO vector (Invitrogen) under the control of the hybrid human cytomegalovirus (CMV)/TetO2 promoter for tetracycline-regulated expression. An HA-tag was added to the C-terminus of UCP1 to facilitate protein detection and normalization. The pcDNA™5/FRT⁄TO vector was chosen for its compatibility with the Flp-In™ T-REx™ system, allowing for site-specific integration and tetracycline-inducible expression. The UCP1 inserts were ligated into pcDNA™5/FRT/TO and transformed into competent TOP10 E. coli (Invitrogen). Transformants were selected on LB agar plates containing 100 μg/ml ampicillin. Colonies were analyzed for the presence and correct orientation of the insert by restriction digestion and Sanger sequencing using CMV Forward and BGH Reverse primers. Cell lines exhibiting tetracycline-inducible (0.3 μg/ml for NMR, 0.2 μg/ml for mouse) expression of UCP1 were generated using Flp-In™ 293 T-REx cells (Invitrogen; R78007) according to the manufacturer’s instructions. Briefly, cells were co-transfected with the pcDNA™5/FRT/TO-UCP1 construct (9 μg) and pOG44 Flp recombinase expression plasmid (1 μg) using Lipofectamine 2000 (Invitrogen) according to the manufacturer’s instructions. Stable integrants were selected using 100 μg/ml hygromycin B (Invitrogen). To induce UCP1 expression, cells were treated with 0.3 μg/ml (NMR) or 0.2 μg/ml (mouse) tetracycline for 24 hours.

Three mutant UCP1 cell lines were generated using site-directed mutagenesis of the mouse UCP1 sequence: (1) G263R+P264W, (2) H146Q, and (3) G263R+P264W+H146Q. These mutations correspond to the NMR-specific amino acid changes in UCP1. Mutagenesis was performed using the QuikChange II Site-Directed Mutagenesis Kit (Agilent) following the manufacturer’s protocol. Primers used for mutagenesis were as follows:

For G263R+P264W:

5’-gaaaaaggccgtccatctttccttggtgtacatggacatcgc-3’
5’-gcgatgtccatgtacaccaaggaaagatggacggcctttttc-3’

For H146Q:

5’-tttgatcccatgcagttggctctgggcttgc-3’
5’-gcaagcccagagccaactgcatgggatcaaa-3’

Successful mutagenesis was confirmed by Sanger sequencing. Mutant constructs were then used to generate stable cell lines as described above.

### Western blotting

Samples of frozen white and brown adipose tissues or cultured cells were lysed in a protein lysis buffer containing protease inhibitors (Roche). Proteins were separated by SDS-PAGE using NuPAGE Novex gels (Invitrogen) and were transferred onto polyvinylidene difluoride membranes (Invitrogen). Blocking was performed in 5% skim milk. The following antibodies were used for Western blot analysis: HA (Abcam, ab9110, 1:1000), UCP1 (Abcam, ab10983, 1:10000), vinculin (Sigma-Aldrich, V9264-200UL, 1:10000), β-actin (Sigma, A3854, 1:10000). Blots were visualized using SuperSignal West Pico PLUS Chemiluminescent Substrate (Thermo Scientific) and ChemiDoc Imaging Systems (Bio-Rad). For UCP1 detection in HEK293 cell lines expressing mouse or NMR UCP1, we used anti-HA antibodies. This approach was chosen to ensure consistent detection between mouse and NMR UCP1, given potential differences in UCP1-antibody affinity between species. It was also used in attempt to more accurately normalize UCP1 content between mouse and NMR UCP1-expressing cell lines based on tetracycline concentration and band density on Western blots. Previous studies have shown that HA-tagged UCP1 generally retains its activity (49, 50), although species-specific effects cannot be ruled out.

### UCP1 localization in HEK 293 cells

Mitochondrial localization was observed by staining live cells with MitoTracker Red CMXRos (Invitrogen, M7512). A 1 mM stock solution of MitoTracker Red CMXRos was prepared in high-quality, anhydrous DMSO according to the manufacturer’s instructions. The stock solution was diluted to a working concentration of 100 nM in prewarmed (37°C) growth medium. Cells grown on coverslips were incubated with the MitoTracker staining solution for 30 minutes under standard growth conditions (37°C, 5% CO2). After staining, cells were washed with fresh, prewarmed growth medium. Cells were then fixed using PBS containing 4% paraformaldehyde (Sigma) for 15 minutes at room temperature. After rinsing with PBS, fixed cells were permeabilized for 10 minutes in PBS containing 0.1% Triton X-100. Following permeabilization, cells were blocked for 1 h in PBS containing 10% (v/v) goat serum. Fixed cells were incubated with primary anti-HA tag antibodies (Abcam, ab9110, 1:1000) overnight at 4 °C. After washing out unbound antibodies, cells were incubated with secondary antibodies (labeled with Alexa 488, 1:200 dilution, Jackson ImmunoResearch) for 1 h at room temperature. Nuclei were counterstained using 1 μg/mL DAPI solution (Life Technologies; 62248, 1mg/mL stock). Fluorescence imaging was conducted using an LSM 700 confocal microscope (Zeiss).

### Cellular oxygen consumption

Approximately 30,000 HEK293 cells expressing mouse and NMR UCP1 were seeded into gelatin-coated XF24 culture microplates (Seahorse Bioscience) and cultured in DMEM (DMEM high glucose + GlutaMAX) with 10% FBS, 1X antibiotic/antimycotic overnight at 37 °C in an atmosphere of 5% CO_2_. The XF24 plates were then transferred to a temperature-controlled (37 °C) XF24 Extracellular Flux analyzer (Seahorse Bioscience). Leak respiration was induced by addition of 1 μM oligomycin (inhibiting ATP synthase) by automatic pneumatic injection. Next, 10 μM of TTNPB (4-[(E)-2-(5,6,7,8-tetrahydro-5,5,8,8-tetramethyl-2-naphthalenyl)-1 propenyl] benzoic acid) diluted in dimethylsulfoxide was injected to specifically activate UCP1, followed by an injection of 0.5 μM FCCP (carbonyl cyanide p trifluoromethoxyphenylhydrazone) to completely uncouple mitochondria. Finally, injection of 2 μM rotenone with 1 μM antimycin A (respiratory chain inhibitors) was performed to measure basal non-mitochondrial respiration.

### Isolation of mitochondria

Mitochondria were isolated from HEK 293T cells and brown adipose tissue (BAT) by homogenization and subsequent differential centrifugation, based on our previously published protocol (51). The extraction buffer contained 10 mM HEPES, 200 mM mannitol, 70 mM sucrose, 1 mM EGTA, and 2 mg/ml BSA, adjusted to pH 7.5. To stabilize thiol-containing metabolites, prevent oxidation of the mitochondrial glutathione pool, and mimic hypoxic conditions during isolation (52, 53), we included 20 mM N-ethylmaleimide (NEM) in the extraction buffer. For HEK293 cells, homogenization was performed using a Dounce homogenizer with a tight pestle (type B), applying 25-40 strokes on ice. This method was chosen to ensure efficient cell membrane disruption while preserving mitochondrial integrity in these softer cells. For BAT, we employed a two-step homogenization process to account for the tissue’s fibrous nature: first, using a loose pestle (type A) for 5-10 strokes to break down larger tissue pieces, followed by a tight pestle (type B) for 10-20 strokes to ensure thorough homogenization. This approach allows for gentle initial disruption of the tissue structure followed by more complete cell lysis. The homogenate was centrifuged at 600 g for 10 min and the supernatant was filtered through 250 μm diameter gauze. It was further centrifuged at 11,000 g for 10 min. The supernatant was collected as the cytosolic fraction. The pellet was resuspended in extraction buffer without BSA and centrifuged at 600 g for 10 min. Then, the supernatant was re-centrifuged at 11,000 g for 10 min. The resulting pellet constituted the mitochondrial fraction. Total mitochondrial protein concentration was quantified using the Bradford method. For membrane potential measurements, 50 μg of mitochondrial protein was used per well. All steps were performed on ice or at 4°C to maintain mitochondrial integrity. This protocol ensures a pure mitochondrial fraction suitable for functional studies, as validated in our previous work (51).

### Mitochondrial respiration and membrane potential measurements

Membrane potential of isolated mitochondria was measured by safranin O fluorescence quenching using a Seahorse XF96 Flux Analyzer (Seahorse Bioscience). This dye accumulates in mitochondria in a membrane potential-dependent manner. As the mitochondrial membrane potential increases (hyperpolarization), more Safranin O accumulates inside the mitochondria, leading to a decrease in fluorescence of the medium. Conversely, as the membrane potential decreases (depolarization), Safranin O is released from the mitochondria, resulting in an increase in fluorescence of the medium. Mitochondria (50 μg protein) were suspended in assay buffer (125 mM sucrose, 20 mM HEPES, 2 mM MgCl2, 1 mM EDTA, pH 7.2) containing 8 μM safranin O. Fluorescence was recorded at excitation and emission wavelengths of 495 nm and 586 nm, respectively, at 37°C. Succinate (5 mM) was used to energize mitochondria and establish a membrane potential. UCP1 was activated by sequential additions of TTNPB (0-30 μM) or inhibited by GDP (0-1 mM). At the end of each experiment, FCCP (2 μM) was injected to achieve complete depolarization and maximum fluorescence (Fmax). For data analysis, we used the following calculations:

- ΔF/Fmax: The change in fluorescence (ΔF) due to TTNPB activation, normalized to the maximum fluorescence (Fmax) achieved with FCCP. Higher values indicate greater UCP1 activation.
- -ΔF/F_TTNPB_: The negative change in fluorescence relative to the TTNPB-activated state, used to quantify GDP inhibition. Higher values indicate greater inhibition.

This normalization allows for comparison between different mitochondrial preparations and UCP1 variants.

### Body Composition

Body composition was examined using the 3-in-1 Echo MRI Composition Analyzer (Echo Medical Systems) prior to temperature probe implantation surgery in mice and NMRs.

### β3-adrenergic receptor activation

For acute monitoring following administration of saline or adrenergic activation (Figure 2*D*, S3*B*), wild type C57Bl/6J mice (Jackson Labs) and naked mole-rats (from the University of Illinois-Chicago colony, 11 months old) were used. Conscious animals were placed into an open-circuit Comprehensive Lab Animal Monitoring System (CLAMS; Columbus Instruments) and maintained at 30 °C for a total of 9 hours. Animals were removed from the cages for less than 60 seconds each to receive an intraperitoneal injection of 200 µl sterile saline solution and returned to the cage for monitoring. Approximately 3 hours following saline administration, the animals were again removed from the cages for less than 60 seconds to receive an intraperitoneal injection of the selective β3-adrenergic receptor agonist CL316,243 (1 mg/kg dissolved in sterile 0.9% NaCl) (Sigma Chemical). Means values were calculated over the entire data collection period following each injection, starting from the first post-injection data point until the end of the monitoring period. At the conclusion of this experiment, animals were housed at 30 °C. A similar injection schedule was followed on the next day, with thermal imaging of the dorsal side of the mouse and NMR without anesthesia or restraint using a FLIR T62101 camera (FLIR Systems, Inc).

### Individual animal housing and acclimation to CLAMS

Naked mole-rats (NMRs) and mice were individually housed in metabolic cages (CLAMS, Columbus Instruments) for the duration of the experiments. While we acknowledge that individual housing can be a source of stress for social animals like NMRs and mice, current technology limits us to this approach for accurate metabolic measurements. To mitigate potential stress, mice were acclimated to the metabolic cages for up to 24 hours prior to data collection. This acclimation period allows for habituation to the new environment, and while it may result in a transient period of increased activity, it does not significantly affect whole-body metabolic rates (54). NMR were not acclimated and did not exhibit increased locomotor activity when individually housed in the indirect calorimetry cages. Throughout the experiments, we monitored several behavioral indicators of stress, including locomotor activity and feeding behavior.

### Body temperature and metabolic measurements during exposure to various ambient temperatures

To evaluate the thermoregulatory state of NMRs, we continuously monitored core body temperature using implanted temperature sensors while exposing animals to a range of ambient temperatures (18-32°C) (Figures 5*A*-*F*, 6*A*-*F*). Additional experiments included challenging non-insulated NMR at 34 °C and 37 °C to establish a more complete ambient temperature challenge response (Fig. 3). NMRs used during exposure to various ambient temperatures, were obtained from established colonies at University of Illinois-Chicago. All subjects (3 males, 2 females) were non-breeding subordinates, approximately 11 months old. NMRs live in eusocial colonies with a strict reproductive hierarchy, where only one female (the queen) and one to three males breed, while the rest remain as non-breeding subordinates. Due to a phenomenon termed ‘socially induced reproductive suppression’ (55), non-breeding subordinate NMRs do not express sex-specific hormones or undergo typical mammalian sexual development. Consequently, subordinate males and females are physiologically very similar and lack the sex-specific hormone profiles that typically drive physiological differences between males and females in other mammals. Given this unique characteristic of NMRs and our relatively small sample size, we did not analyze sex differences in our study, as all subjects were expected to be physiologically similar regardless of genetic sex.

Mice were shaved using electric clippers while being anesthetized with 1-2% isoflurane. Both those with intact fur (n=9) and shaved mice (n=9) were simultaneously housed individually in the CLAMS. These 10-week-old male mice were fed Picolab Rodent Diet 5053 (LabDiet) Chow, and NMRs were on fresh raw yams, apples, and bananas, as well as on a vitamin and protein enriched cereal (Pronutro, Pioneer Foods). All animals were maintained with a 12-hour light and dark cycle (07:00-19:00). Mice were acclimated for one day prior to the experiment. NMRs were not acclimated to maintain their group housing for as long as possible to avoid undue stress. Rates of O_2_ consumption (V_O2_) and CO_2_ production (V_CO2_), as well as core body temperatures were experimentally determined. Respiratory Exchange Ratio (RER) was calculated as the ratio of CO2 production to O2 consumption (RER = VCO2/VO2). This measure provides insight into substrate utilization, with lower values consistent with fat oxidation and higher values indicating primarily carbohydrate oxidation. Rates of energy expenditure (kcal/h) were calculated from gas exchange. For these experiments, animals were maintained at 30 °C for mice and 32 °C for NMRs. Animals were challenged with 24-hour exposure to each of the subsequent temperatures followed by 24-48 hours to return to thermoneutrality. NMR were challenged with 28 °C, 25 °C, 22 °C, and 18 °C (the latter only for 12 hours). Mice were challenged with 28 °C, 25 °C, 22 °C, 18 °C, 14 °C and 6 °C. The temperature transition from thermoneutrality occurred over three hours at a continuous rate and was excluded from analysis to focus on physiological responses during stable temperature periods. Calorimetry data, collected every 12 minutes throughout the experiments, were binned per hour and outliers removed with CalR (56).

### Thermogenesis following olive oil supplementation

In the olive oil supplementation experiment (Figure 4*A*), extra Virgin Olive Oil was administered by gavage (5 μl/g body weight) to animals maintained at 30 °C (N=5 mice) and 32°C (N=5 NMRs). Cages were transitioned to a cold challenge over 3 hours at a constant rate, mice from 30 °C to 4 °C (57) and NMR from 32°C to 22°C. Data were analyzed from the entire stable temperature period at 22°C (NMRs), following the 3-hour transition from 32°C. Animals were not fasted and had free access to food throughout the experiment. We did not perform blood draws to measure circulating non-esterified fatty acid (NEFA) levels after gavage or during cold exposure. This decision was made to minimize stress and avoid potential confounding effects on catecholamine release, lipolysis, and thermogenesis, which could be triggered by the handling required for blood collection in naked mole-rats. We assumed that the bolus administration of fatty acids would increase circulating fatty acid levels in NMRs in a manner analogous to that observed in mice and other rodents. We acknowledge that the lack of direct NEFA measurements is a limitation of our study.

### Insulation experiments

NMR insulation was provided by introducing fleece Pet Beds (Petco, Inc.) tied at the top to create an enclosure (Fig. 6*G*). This approach was chosen after initial attempts to provide constant insulation using custom-made wool sweaters proved unsuccessful, as NMRs showed signs of distress and actively tried to remove the garments (Fig. S4). The fleece bed method allowed NMRs to enter and leave the insulated area voluntarily, mimicking their natural burrowing behavior. As a result, our reported average body temperatures and energy expenditures (Fig. 6*A-F*) represent a mix of in-shelter and out-of-shelter periods. While this approach introduced some variability in insulation exposure (Fig. S5), it minimized stress on the animals and better reflected their natural thermoregulatory behaviors.

### Temperature Probe Implantation Surgery

Both mice and NMRs were surgically implanted with wireless temperature probes (G2 E-Mitter, Starr Lifescience) in the peritoneal cavity. Both groups of animals were anesthetized with 1-2% isoflurane until non-responsive to a toe pinch. Animals were administered with 0.1 mg/kg Buprenorphine subcutaneously before surgery. A 2 cm midline abdominal skin incision was made 1 cm below the diaphragm. The abdominal cavity was then opened by making a 2 cm. incision along the posterior left lower quadrant. Temperature probes were implanted into the peritoneal cavity and secured with non-absorbable 4-0 sutures. The abdominal cavity was closed using an absorbable 4-0 suture and the skin was closed with monofilament non-absorbable 4-0 sutures using a continuous running stitch. Pain relief was provided every 12 hours for 48 hours postoperatively with 0.1 mg/kg Buprenorphine subcutaneously. All animals were given 7 days to recover while housed at 30-32 °C prior to experimentation.

### Thermoneutral temperature determination

The thermoneutral temperature for NMRs (Fig. 3*B*) was determined using the Scholander plot approach (58). We measured metabolic rates of NMRs at various ambient temperatures ranging from 18°C to 37°C. The thermoneutral temperature was identified as the ambient temperature at which metabolic rate was lowest and began to plateau, indicating the point at which no additional energy expenditure was required for thermoregulation.

### Statistical analysis

All data are presented as mean values ± SD or ±SEM. For comparison of the variation within the tested groups to the variation among the means of the groups, F-test was performed. One-way repeated measures ANOVA was performed to evaluate the effect of T_a_ on dependent variables (T_b_, EE or RER). A two-way repeated measures ANOVA (for NMRs) and two-way mixed ANOVA (for mice) was performed to evaluate the effect of insulation over various T_a_ on animals’ T_b_. Tukey’s test or paired t-test with the Bonferroni multiple testing correction method were used for pairwise comparisons of dependent variables between T_a_ groups at each insulation group. Normality assumption was assessed by Shapiro-Wilk’s test. Assumption of sphericity was tested using Mauchly’s Test for Sphericity. Greenhouse-Geisser sphericity correction was applied to factors violating the sphericity assumption. Homogeneity of variances was assessed using Levene’s test for equality of variances. Box’s M-test was used to evaluate the homogeneity of covariances for shaved vs furry groups of mice. p < 0.05 was considered statistically significant. See Tables S1-S4 for more details.

## Acknowledgments

Supported by grants from the National Institute on Aging.

## Author Contributions

V.N.G. and A.S.B. conceived the study. V.N.G., A.S.B., and A.M. designed experiments. A.M., J.C., Y.H., Z.P., S.H.Y, S.L., Z.D., V.G.A., A.S.B. performed experiments. A.M., J.C., and A.S.B. analyzed and interpreted the data. V.N.G. and A.S.B. supervised the study. V.G., A.S., and T.J.P provided advice about experimental design. T.J.P provided naked mole-rats. A.M., V.N.G., and A.S.B. wrote the manuscript with a contribution from S.H.Y, S.L., V.G., and T.J.P.

