## Supplemental materials for "Comparative analysis of naked mole-rat thermogenesis and its potential to maintain euthermia in response to cold"

Supplementary Information
for

### Comparative analysis of naked mole-rat thermogenesis and its potential to maintain euthermia in response to cold

Aleksei Mikhalchenko[a], June K. Corrigan[b], Yuchen He[b], Zalan Peterfi[a], Sun Hee Yim[a,1], Sang-Goo Lee[a], Zhaoming Deng[b], Vince G. Amoroso[c], Vera Gorbunova[d], Andrei Seluanov[d], Thomas J. Park[c], Alexander S. Banks[b,*], Vadim N. Gladyshev[a,*]

[a] Division of Genetics, Department of Medicine, Brigham and Women's Hospital, Harvard Medical School, Boston, MA 02115, USA

[b] Division of Endocrinology, Diabetes and Metabolism, Beth Israel Deaconess Medical Center, Harvard Medical School, Boston, MA 02215, USA

[c] Department of Biological Sciences, University of Illinois at Chicago, Chicago, IL 60607, USA
[d] University of Rochester, Department of Biology, Rochester, NY 14627, USA

[1] Present address: The Institute of Environmental and Human Health, Texas Tech University, Lubbock, TX 79409

**Keywords:** naked mole-rat, thermogenesis, UCP1, brown adipose tissue, metabolism

**This includes:**

Figures S1 to S5
Tables S1 to S4

```
NMR    MVSPTASEVHPTMGVKIFSAGVAACLADVITFPLDTAKVRLQIQGESQISSGIQYKGALG 60
DMR    MGSPTASEVHPTVGVKIFSAGVSACLADVITFPLDTAKVRLQIQGESQISSGIRYKGVLG 60
Guinea MVSPTVSDVHPTMGVKIFSAGVSACLADVITFPLDTAKVRLQIQGESPTSSGIRYKGVLG 60
Mouse  MVNPTTSEVQPTMGVKIFSAGVSACLADIITFPLDTAKVRLQIQGEGQASSTIRYKGVLG 60
Rat    MVSSTTSEVQPTMGVKIFSAGVSACLADIITFPLDTAKVRLQIQGEGQASSTIRYKGVLG 60
Cat    MVGLTASDVPPTMAVKIFSAGVAACVADVITFPLDTAKVRLQIQGECQTSSTIKYRGVLG 60
Human  MGGLTASDVHPTLGVQLFSAGIAACLADVITFPLDTAKVRLQVQGECPTSSVIRYKGVLG 60

NMR    TITTLAKTEGPMKLYSGLPAGLQRQISFASLRIGLYDTVQEFYTSEKDITPSLGSRIAAG 120
DMR    TITTLAKTEGPVKLYSGLPAGLQRQISFASLRIGLYDTVQEFYTSEKDTTPSLGNRIAAG 120
Guinea TITTLAKTEGPVKLYSGLPAGLQRQISFASLRIGLYDTVQEFYTSEKDATPSLGSRIAAG 120
Mouse  TITTLAKTEGLPKLYSGLPAGIQRQISFASLRIGLYDSVQEYFSSGRETPASLGNKISAG 120
Rat    TITTLAKTEGLPKLYSGLPAGIQRQISFASLRIGLYDTVQEYFSSGRETPASLGSKISAG 120
Cat    TITTLAKTEGPMKLYSGLPAGLQRQISFASLRIGLYDTVQEFFSAGKETTAGLGSKISAG 120
Human  TITAVVKTEGRMKLYSGLPAGLQRQISSASLRIGLYDTVQEFLTAGKETAPSLGSKILAG 120

NMR    LTTGGVAVFIGQPTEVVKVRLQAQSQLHGLKPRYTGTYNAYRIIATTESLKSLWKGTTPN 180
DMR    LTTGGVAVFIGQPTEVVKVRLQAQSHLHGLKPRYTGTYNAYRIIATTESLKSLWKGTTPN 180
Guinea LTTGGVAVFIGQPTEVVKVRLQAQSHLHGLKPRYTGTYNAYRIIATTESLKSLWKGTTPN 180
Mouse  LMTGGVAVFIGQPTEVVKVRMQAQSHLHGIKPRYTGTYNAYRVIATTESLSTLWKGTTPN 180
Rat    LMTGGVAVFIGQPTEVVKVRMQAQSHLHGIKPRYTGTYNAYRVIATTESLSTLWKGTTPN 180
Cat    LTTGGVAVFIGQPTEVVKVRLQAQSHLHGLKPRYTGTYNAYRIIATTEGLTGLWKGTTPN 180
Human  LTTGGVAVFIGQPTEVVKVRLQAQSHLHGIKPRYTGTYNAYRIIATTEGLTGLWKGTTPN 180

NMR    LLRNIIINCTELVTYDLMKGALVRNKILADDVPCHLLSALIAGFCTTLLSSPVDVVKTRF 240
DMR    LLRNIIINCTELVTYDLMKGALVRNKILADDIPCHLLSALIAGFCTTLLSSPVDVVKTRF 240
Guinea LLRNIIINCTELVTYDLMKGALVRNKILADDVPCHLLSALIAGFCTTLLSSPVDVVKTRF 240
Mouse  LMRNVIINCTELVTYDLMKGALVNNKILADDVPCHLLSALVAGFCTTLLASPVDVVKTRF 240
Rat    LMRNVIINCTELVTYDLMKGALVNHHILADDVPCHLLSALVAGFCTTLLASPVDVVKTRF 240
Cat    LTRNVIINCTEIVTYDLMKEALVKNKLLADDLPCHFVSALIAGFCTTVLSSPVDVVKTRF 240
Human  LMRSVIINCTELVTYDLMKEAFVKNNILADDVPCHLVSALIAGFCATAMSSPVDVVKTRF 240

NMR    INSPPGQYISVPSCAMTMLKKERWTAFFKGFVPSFLRLASWNVIMFVCFEQLKRELTKSR 300
DMR    INSAPGQYVSVPSCAMTMLNKEGPTAFFKGFVPSFLRLASWNVIMFVCFEQLKRELTKSG 300
Guinea INSPPGQYLSVPSCAMTMLLKEGPTAFFKGFVPSFLRLASWNVIMFVCFEQLKRELMKSR 300
Mouse  INSLPGQYPSVPSCAMSMYTKEGPTAFFKGFVASFLRLGSWNVIMFVCFEQLKKELMKSR 300
Rat    INSLPGQYPSVPSCAMTMYTKEGPAAFFKGFAPSFLRLGSWNVIMFVCFEQLKKELMKSR 300
Cat    VNSPPGQYTSVPNCAITMLTKEGPLAFFKGFVPSFLRLGSWNVIMFVCFEQLKRELMKSR 300
Human  INSPPGQYKSVPNCAMKVFTNEGPTAFFKGLVPSFLRLGSWNVIMFVCFEQLKRELSKSR 300

NMR    QPTDYAT 307
DMR    QSADYAA 307
Guinea QPVDYAT 307
Mouse  QTVDCTT 307
Rat    QTVDCTT 307
Cat    QTMDCAT 307
Human  QTMDCAT 307
```

**Fig. S1. Unique sequence changes in NMR UCP1.** Alignment of mammalian UCP1 sequences. Species shown are: Homo sapiens (Human), Mus musculus (Mouse), Rattus norvegicus (Rat), Heterocephalus glaber (Naked mole-rat, NMR), Fukomys damarensis (Damaraland mole-rat, DMR), Cavia porcellus (Guinea pig), Felis catus (Cat). Amino acids unique to the NMR are highlighted in red. Conserved motifs, highlighted in blue, represent functional domains crucial for UCP1 activity, including nucleotide binding sites and regions involved in proton transport. These conserved motifs underscore the functional importance of these regions across diverse mammalian species. Identical residues are shaded in black. The three amino acid changes specific to NMR UCP1 are: H146Q, G263R, P264W.

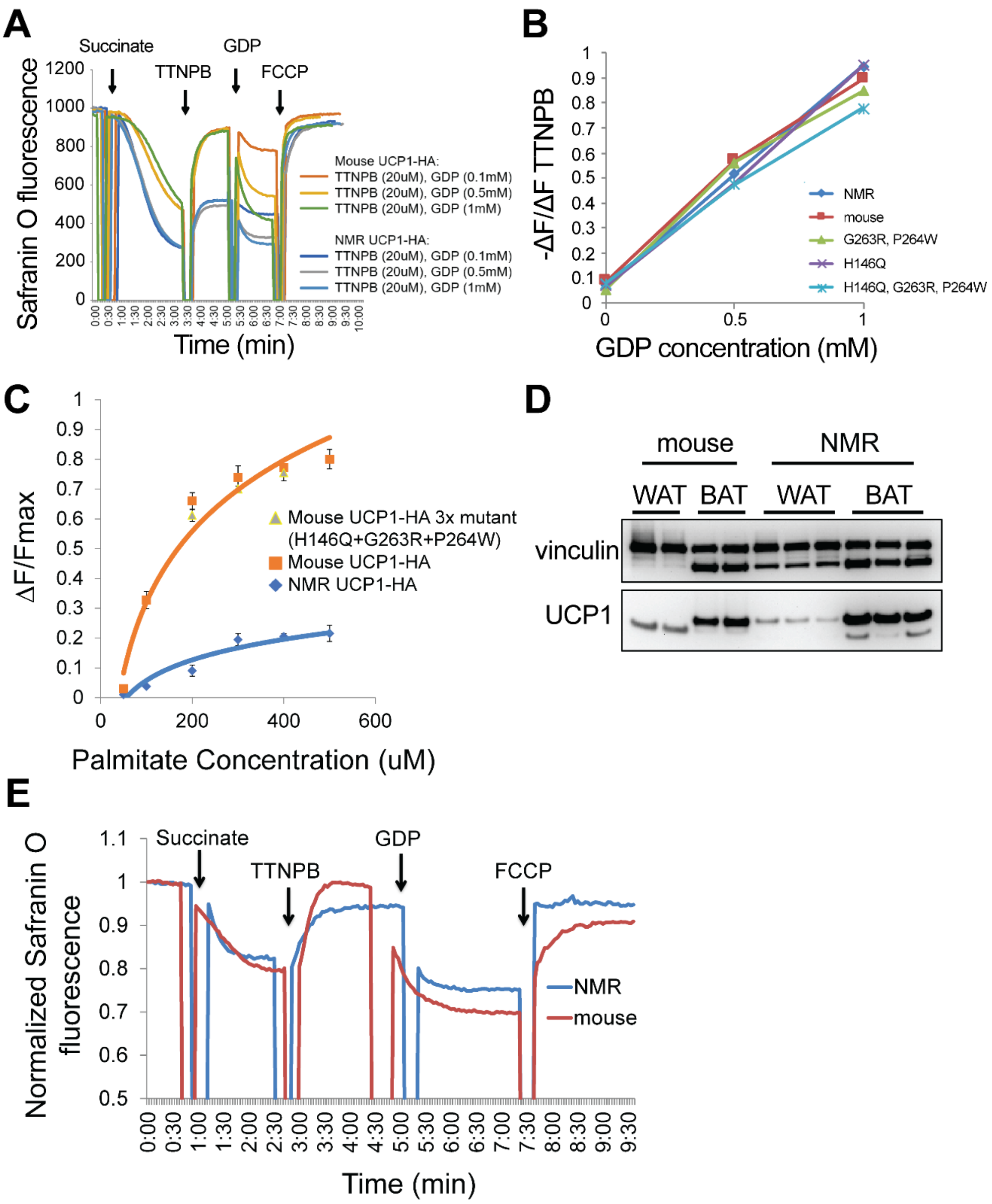

**Fig. S2. Characterization of NMR BAT and UCP1 function *in vitro*. Related to Figure 2.**

(*A*) Real-time changes in safranin O fluorescence in mitochondria isolated from HEK293 cells expressing NMR and mouse UCP1-HA. Safranin O fluorescence was measured upon sequential addition of succinate (substrate), TTNPB (UCP1 activator), GDP (UCP1 inhibitor), and FCCP (to achieve maximum depolarization). Decreases in fluorescence indicate mitochondrial polarization, while increases indicate depolarization. NMR UCP1 shows a lower response to TTNPB activation compared to mouse UCP1. (*B*) GDP inhibition of UCP1 activity in mitochondria from HEK293 cells expressing NMR, mouse, and mutant mouse UCP1 forms. The y-axis ($-\Delta F/F_{TTNPB}$) represents the negative change in fluorescence relative to the TTNPB-activated state, with higher values indicating greater inhibition by GDP. GDP causes repolarization (decrease in fluorescence) from the depolarized state induced by

TTNPB. Similar slopes for all UCP1 variants suggest comparable GDP inhibition. (*C*) Relative change in mitochondrial membrane potential ($\Delta F/F_{max}$) in HEK293 cells expressing NMR, mouse, and mouse triple mutant (H146Q, G263R, P264W) UCP1 with palmitate as UCP1 activator. Higher $\Delta F/F_{max}$ values indicate greater depolarization, reflecting increased UCP1 activity. NMR UCP1 shows lower maximal activation compared to mouse UCP1 and its mutant forms. All values are mean ± SD from three independent experiments. (*D*) Western blot analysis of NMR and mouse UCP1 in the brown (BAT) and white adipose (WAT) tissues of these animals. Vinculin expression was used as a loading control. (*E*) Real-time changes in safranin O fluorescence in mitochondria isolated from NMR and mouse BAT. Safranin O fluorescence was measured upon sequential addition of succinate (substrate), 10uM TTNPB (UCP1 activator), 1mM GDP (UCP1 inhibitor), and FCCP (to achieve maximum depolarization). Increases in fluorescence indicate depolarization (activation of UCP1 by TTNPB), while decreases indicate repolarization (inhibition of UCP1 by GDP).

A

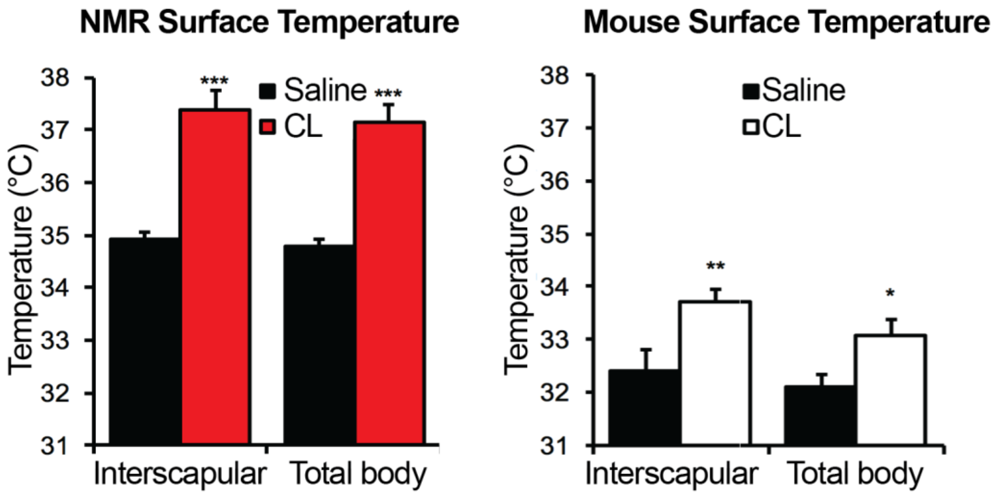

B

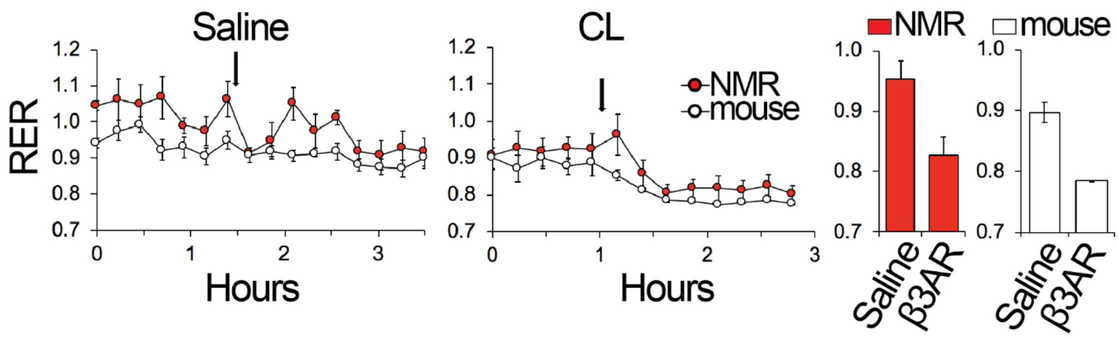

**Fig. S3. Response of NMRs and mice to β3-adrenergic agonist CL-316,243 (CL) interventions. Related to Figure 2.**

(*A*) NMR and mouse average surface temperature (* $p<0.05$, ** $p<0.01$, *** $p<0.001$). (*B*) NMR and mouse respiratory exchange ratio (RER) (all bar plots reach statistical significance ($p<0.05$)). All values are mean ± SEM from five mice and four NMRs.

**Table S1. The relationship between ambient temperature and core body temperature in individual non-anesthetized NMRs. Related to Figure 3.**

| **Effect of $T_a$ on NMR $T_b$ One-way repeated measures ANOVA (type III tests)** | | | | | |
|---|---|---|---|---|---|
| Effect | DFn | DFd | F | p | ges |
| $T_a$ | 6 | 24 | 436.25 | 2.83e-23 | 0.988 |

| **Tukey's test shows pairwise comparisons of NMR $T_b$ between all $T_a$ groups** | | |
|---|---|---|
| **$T_a$** | **$T_b$ difference** | **p** |
| 22-18 | 4.64 | <.0001 |
| 25-18 | 7.14 | <.0001 |
| 28-18 | 9.83 | <.0001 |
| 32-18 | 11.80 | <.0001 |
| 34-18 | 13.04 | <.0001 |
| 37-18 | 15.28 | <.0001 |
| 25-22 | 2.50 | <.0001 |
| 28-22 | 5.18 | <.0001 |
| 32-22 | 7.16 | <.0001 |
| 34-22 | 8.40 | <.0001 |
| 37-22 | 10.64 | <.0001 |
| 28-25 | 2.68 | <.0001 |
| 32-25 | 4.66 | <.0001 |
| 34-25 | 5.90 | <.0001 |
| 37-25 | 8.14 | <.0001 |
| 32-28 | 1.98 | 0.0002 |
| 34-28 | 3.22 | <.0001 |
| 37-28 | 5.46 | <.0001 |
| 34-32 | 1.24 | 0.0285 |
| 37-32 | 3.48 | <.0001 |
| 37-34 | 2.24 | <.0001 |

| **$T_a$** | **Mean $T_b$** | **SD** | **N** |
|---|---|---|---|
| 18 | 21.32 | 1.040 | 5 |
| 22 | 25.96 | 0.434 | 5 |
| 25 | 28.46 | 0.654 | 5 |
| 28 | 31.14 | 0.796 | 5 |
| 32 | 33.12 | 0.383 | 5 |
| 34 | 34.36 | 0.230 | 5 |
| 37 | 36.60 | 0.316 | 5 |

**Table S2. Substrate availability limits thermogenesis in NMRs and mice. Related to Figure 4.**

| Paired t-test | | | |
|---|---|---|---|
| **mean of the differences** | | | **p** |
| 2.62 | | | 0.012 |
| **NMR group** | **Mean $T_b$** | **SD** | **N** |
| Control | 24.88 | 0.976 | 5 |
| Olive oil | 27.50 | 0.374 | 5 |
| **Paired t-test** | | | |
| **mean of the differences** | | | **p** |
| 0.074 | | | 0.639 |
| **mouse group** | **Mean $T_b$** | **SD** | **N** |
| Control | 34.25 | 0.496 | 5 |
| Olive oil | 34.32 | 0.766 | 5 |

**Table S3. Role of insulation in metabolism and thermoregulation of mice. Related to Figure 5.**

| Effect of insulation across tested $T_a$ groups on mouse $T_b$ Two-way mixed ANOVA (type II tests) | | | | | |
|---|---|---|---|---|---|
| **Effect** | **DFn** | **DFd** | **F** | **p** | **ges** |
| insulation | 1 | 16 | 31.321 | 4.01e-05 | 0.38 |
| $T_a$ | 7 | 112 | 37.173 | 1.97e-26 | 0.62 |
| Insulation : $T_a$ | 7 | 112 | 3.507 | 0.002 | 0.13 |

| Effect of insulation at each $T_a$ group Pairwise comparisons of mean $T_b$ between shaved and furry mice (t-test, Bonferroni adj.) | | | | |
|---|---|---|---|---|
| **$T_a$** | **group 1** | **group 2** | **p.adj** | **p.adj.signif** |
| 6 | furry | shaved | 0.0000128 | **** |
| 10 | furry | shaved | 0.0000753 | **** |
| 14 | furry | shaved | 0.0242 | * |
| 18 | furry | shaved | 0.04 | * |
| 22 | furry | shaved | 0.000748 | *** |
| 25 | furry | shaved | 0.0268 | * |
| 28 | furry | shaved | 0.419 | ns |
| 30 | furry | shaved | 0.0117 | * |

| Effect of $T_a$ on mouse $T_b$ One-way repeated measures ANOVA (type III tests) | | | | |
|---|---|---|---|---|
| **insulation** | **Effect** | **F** | **p** | **ges** |
| furry | $T_a$ | 13.37 | 3.07e-05 | 0.56 |
| shaved | $T_a$ | 24.93 | 4.63e-15 | 0.67 |

| Pairwise comparisons of mean $T_b$ in shaved and furry mice (paired t-test, Bonferroni adj.) | | | | |
|---|---|---|---|---|
| **insulation** | **group 1** | **group 2** | **p.adj** | **p.adj.signif** |
| furry | 6 | 10 | 1 | ns |
| furry | 6 | 14 | 1 | ns |
| furry | 6 | 18 | 0.588 | ns |
| furry | 6 | 22 | 0.005 | ** |
| furry | 6 | 25 | 0.025 | * |
| furry | 6 | 28 | 0.088 | ns |
| furry | 6 | 30 | 0.003 | ** |
| furry | 10 | 14 | 1 | ns |
| furry | 10 | 18 | 1 | ns |
| furry | 10 | 22 | 0.005 | ** |
| furry | 10 | 25 | 0.078 | **ns** |
| furry | 10 | 28 | 0.029 | * |
| furry | 10 | 30 | 0.00049 | *** |
| furry | 14 | 18 | 0.037 | * |
| furry | 14 | 22 | 0.029 | * |
| furry | 14 | 25 | 0.0003 | *** |
| furry | 14 | 28 | 0.087 | ns |
| furry | 14 | 30 | 0.041 | * |
| furry | 18 | 22 | 1 | **ns** |
| furry | 18 | 25 | 0.305 | ns |
| furry | 18 | 28 | 1 | ns |
| furry | 18 | 30 | 1 | ns |
| furry | 22 | 25 | 1 | ns |
| furry | 22 | 28 | 1 | ns |
| furry | 22 | 30 | 1 | ns |
| furry | 25 | 28 | 1 | ns |
| furry | 25 | 30 | 1 | ns |
| furry | 28 | 30 | 1 | ns |
| shaved | 6 | 10 | 0.574 | ns |
| shaved | 6 | 14 | 0.251 | ns |
| shaved | 6 | 18 | 0.003 | ** |
| shaved | 6 | 22 | 0.00059 | *** |

| | | | | |
|---|---|---|---|---|
| shaved | 6 | 25 | 0.011 | * |
| shaved | 6 | 28 | 0.00001 | **** |
| shaved | 6 | 30 | 0.00001 | **** |
| shaved | 10 | 14 | 1 | ns |
| shaved | 10 | 18 | 0.375 | ns |
| shaved | 10 | 22 | 0.085 | ns |
| shaved | 10 | 25 | 0.042 | * |
| shaved | 10 | 28 | 0.000068 | **** |
| shaved | 10 | 30 | 0.003 | ** |
| shaved | 14 | 18 | 0.198 | ns |
| shaved | 14 | 22 | 0.154 | ns |
| shaved | 14 | 25 | 0.538 | ns |
| shaved | 14 | 28 | 0.007 | ** |
| shaved | 14 | 30 | 0.015 | * |
| shaved | 18 | 22 | 1 | ns |
| shaved | 18 | 25 | 1 | ns |
| shaved | 18 | 28 | 0.116 | ns |
| shaved | 18 | 30 | 0.288 | ns |
| shaved | 22 | 25 | 1 | ns |
| shaved | 22 | 28 | 0.012 | * |
| shaved | 22 | 30 | 0.605 | ns |
| shaved | 25 | 28 | 0.703 | ns |
| shaved | 25 | 30 | 1 | ns |
| shaved | 28 | 30 | 1 | ns |

| $T_a$ | mouse group | Mean $T_b$ | SD | N |
|---|---|---|---|---|
| 6 | furry | 35.5 | 0.529 | 9 |
| 6 | shaved | 34.2 | 0.360 | 9 |
| 10 | furry | 35.9 | 0.206 | 9 |
| 10 | shaved | 34.7 | 0.644 | 9 |
| 14 | furry | 35.7 | 0.557 | 9 |
| 14 | shaved | 34.9 | 0.787 | 9 |
| 18 | furry | 36.2 | 0.583 | 9 |
| 18 | shaved | 35.6 | 0.577 | 9 |
| 22 | furry | 36.6 | 0.353 | 9 |
| 22 | shaved | 35.7 | 0.556 | 9 |
| 25 | furry | 36.7 | 0.451 | 9 |
| 25 | shaved | 35.98 | 0.766 | 9 |
| 28 | furry | 36.67 | 0.367 | 9 |
| 28 | shaved | 36.5 | 0.477 | 9 |
| 30 | furry | 36.7 | 0.173 | 9 |
| 30 | shaved | 36.4 | 0.265 | 9 |

**Effect of insulation across tested $T_a$ groups on mouse EE**
**Two-way mixed ANOVA (type II tests)**

| Effect | DFn | DFd | F | p | ges |
|---|---|---|---|---|---|
| insulation | 1 | 16 | 32.123 | 3.50e-05 | 0.46 |
| $T_a$ | 7 | 112 | 356.67 | 1.87e-73 | 0.93 |
| Insulation : $T_a$ | 7 | 112 | 11.614 | 4.96e-11 | 0.29 |

**Effect of insulation at each $T_a$ group**
**Pairwise comparisons of mean $T_b$ between shaved and furry mice (t-test, Bonferroni adj.)**

| EE | group 1 | group 2 | p.adj | p.adj.signif |
|---|---|---|---|---|
| 6 | furry | shaved | 0.00000469 | **** |
| 10 | furry | shaved | 0.00000004 | **** |
| 14 | furry | shaved | 0.0000935 | **** |
| 18 | furry | shaved | 0.00933 | ** |

| | | | | |
|---|---|---|---|---|
| 22 | furry | shaved | 0.00609 | ** |
| 25 | furry | shaved | 0.277 | ns |
| 28 | furry | shaved | 0.217 | ns |
| 30 | furry | shaved | 0.724 | ns |

**Effect of $T_a$ on mouse EE**
**One-way repeated measures ANOVA (type III tests)**

| insulation | Effect | F | p | ges |
|---|---|---|---|---|
| furry | $T_a$ | 210.5 | 1.92e-12 | 0.93 |
| shaved | $T_a$ | 173.33 | 1.46e-35 | 0.93 |

**Pairwise comparisons of mean EE in shaved and furry mice (paired t-test, Bonferroni adj.)**

| insulation | group 1 | group 2 | p.adj | p.adj.signif |
|---|---|---|---|---|
| furry | 6 | 10 | 4.00e-03 | ** |
| furry | 6 | 14 | 6.00e-03 | ** |
| furry | 6 | 18 | 6.47e-05 | **** |
| furry | 6 | 22 | 2.60e-08 | **** |
| furry | 6 | 25 | 4.45e-08 | **** |
| furry | 6 | 28 | 4.40e-07 | **** |
| furry | 6 | 30 | 4.93e-08 | **** |
| furry | 10 | 14 | 2.10e-02 | * |
| furry | 10 | 18 | 2.48e-05 | **** |
| furry | 10 | 22 | 4.26e-07 | **** |
| furry | 10 | 25 | 2.20e-08 | **** |
| furry | 10 | 28 | 5.21e-07 | **** |
| furry | 10 | 30 | 3.89e-08 | **** |
| furry | 14 | 18 | 6.00e-03 | ** |
| furry | 14 | 22 | 1.10e-02 | * |
| furry | 14 | 25 | 2.00e-04 | *** |
| furry | 14 | 28 | 1.64e-04 | *** |
| furry | 14 | 30 | 2.54e-04 | *** |
| furry | 18 | 22 | 5.80e-02 | ns |
| furry | 18 | 25 | 9.49e-05 | **** |
| furry | 18 | 28 | 6.13e-05 | **** |
| furry | 18 | 30 | 1.27e-04 | *** |
| furry | 22 | 25 | 1.35e-04 | *** |
| furry | 22 | 28 | 5.91e-05 | **** |
| furry | 22 | 30 | 2.15e-05 | **** |
| furry | 25 | 28 | 3.00e-03 | ** |
| furry | 25 | 30 | 3.30e-02 | * |
| furry | 28 | 30 | 1.23e-01 | ns |
| shaved | 6 | 10 | 1.19e-01 | ns |
| shaved | 6 | 14 | 3.00e-03 | ** |
| shaved | 6 | 18 | 1.31e-04 | *** |
| shaved | 6 | 22 | 8.40e-06 | **** |
| shaved | 6 | 25 | 3.16e-06 | **** |
| shaved | 6 | 28 | 5.18e-07 | **** |
| shaved | 6 | 30 | 1.35e-09 | **** |
| shaved | 10 | 14 | 1.10e-02 | * |
| shaved | 10 | 18 | 3.50e-04 | *** |
| shaved | 10 | 22 | 1.07e-05 | **** |
| shaved | 10 | 25 | 1.13e-06 | **** |
| shaved | 10 | 28 | 2.24e-07 | **** |
| shaved | 10 | 30 | 8.93e-10 | **** |
| shaved | 14 | 18 | 6.60e-02 | ns |
| shaved | 14 | 22 | 2.00e-03 | ** |
| shaved | 14 | 25 | 2.69e-04 | *** |
| shaved | 14 | 28 | 4.62e-05 | **** |
| shaved | 14 | 30 | 1.02e-07 | **** |
| shaved | 18 | 22 | 7.20e-02 | ns |

| | | | | |
|---|---|---|---|---|
| shaved | 18 | 25 | 6.00e-03 | ** |
| shaved | 18 | 28 | 2.83e-04 | *** |
| shaved | 18 | 30 | 1.43e-04 | *** |
| shaved | 22 | 25 | 4.80e-02 | * |
| shaved | 22 | 28 | 2.65e-04 | *** |
| shaved | 22 | 30 | 5.77e-04 | *** |
| shaved | 25 | 28 | 3.70e-02 | * |
| shaved | 25 | 30 | 1 | ns |
| shaved | 28 | 30 | 1 | ns |

| $T_a$ | mouse group | Mean EE | SD | N |
|---|---|---|---|---|
| 6 | furry | 0.809 | 0.05 | 9 |
| 6 | shaved | 0.985 | 0.06 | 9 |
| 10 | furry | 0.744 | 0.047 | 9 |
| 10 | shaved | 0.943 | 0.039 | 9 |
| 14 | furry | 0.638 | 0.082 | 9 |
| 14 | shaved | 0.826 | 0.072 | 9 |
| 18 | furry | 0.566 | 0.055 | 9 |
| 18 | shaved | 0.681 | 0.103 | 9 |
| 22 | furry | 0.486 | 0.029 | 9 |
| 22 | shaved | 0.577 | 0.082 | 9 |
| 25 | furry | 0.412 | 0.035 | 9 |
| 25 | shaved | 0.438 | 0.061 | 9 |
| 28 | furry | 0.324 | 0.041 | 9 |
| 28 | shaved | 0.353 | 0.054 | 9 |
| 30 | furry | 0.361 | 0.036 | 9 |
| 30 | shaved | 0.369 | 0.056 | 9 |

**Effect of insulation across tested $T_a$ groups on mouse RER**
**Two-way mixed ANOVA (type II tests)**

| Effect | DFn | DFd | F | p | ges |
|---|---|---|---|---|---|
| insulation | 1 | 16 | 4.63 | 4.70e-02 | 0.025 |
| $T_a$ | 7 | 112 | 31.61 | 8.44e-24 | 0.64 |
| Insulation : $T_a$ | 7 | 112 | 4.4 | 2.35e-04 | 0.2 |

**Effect of insulation at each $T_a$ group**
**Pairwise comparisons of mean $T_b$ between shaved and furry mice (t-test, Bonferroni adj.)**

| RER | group 1 | group 2 | p.adj | p.adj.signif |
|---|---|---|---|---|
| 6 | furry | shaved | 0.439 | ns |
| 10 | furry | shaved | 0.613 | ns |
| 14 | furry | shaved | 0.432 | ns |
| 18 | furry | shaved | 0.134 | ns |
| 22 | furry | shaved | 0.872 | ns |
| 25 | furry | shaved | 0.05 | ns |
| 28 | furry | shaved | 0.109 | ns |
| 30 | furry | shaved | 0.00246 | ** |

**Effect of $T_a$ on mouse RER**
**One-way repeated measures ANOVA (type III tests)**

| insulation | Effect | F | p | ges |
|---|---|---|---|---|
| furry | $T_a$ | 9.53 | 0.002 | 0.53 |
| shaved | $T_a$ | 25.9 | 1.08e-07 | 0.74 |

**Pairwise comparisons of mean RER in shaved and furry mice (paired t-test, Bonferroni adj.)**

| insulation | group 1 | group 2 | p.adj | p.adj.signif |
|---|---|---|---|---|
| furry | 6 | 10 | 1 | ns |
| furry | 6 | 14 | 1 | ns |

| | | | | |
|---|---|---|---|---|
| furry | 6 | 18 | 1 | ns |
| furry | 6 | 22 | 0.215 | ns |
| furry | 6 | 25 | 0.969 | ns |
| furry | 6 | 28 | 0.006 | ** |
| furry | 6 | 30 | 0.002 | ** |
| furry | 10 | 14 | 1 | ns |
| furry | 10 | 18 | 1 | ns |
| furry | 10 | 22 | 1 | ns |
| furry | 10 | 25 | 1 | ns |
| furry | 10 | 28 | 1 | ns |
| furry | 10 | 30 | 0.00086 | *** |
| furry | 14 | 18 | 1 | ns |
| furry | 14 | 22 | 1 | ns |
| furry | 14 | 25 | 0.073 | ns |
| furry | 14 | 28 | 0.101 | ns |
| furry | 14 | 30 | 0.057 | ns |
| furry | 18 | 22 | 1 | ns |
| furry | 18 | 25 | 1 | ns |
| furry | 18 | 28 | 1 | ns |
| furry | 18 | 30 | 0.153 | ns |
| furry | 22 | 25 | 1 | ns |
| furry | 22 | 28 | 1 | ns |
| furry | 22 | 30 | 0.093 | ns |
| furry | 25 | 28 | 1 | ns |
| furry | 25 | 30 | 1 | ns |
| furry | 28 | 30 | 0.128 | ns |
| shaved | 6 | 10 | 0.207 | ns |
| shaved | 6 | 14 | 1 | ns |
| shaved | 6 | 18 | 0.044 | * |
| shaved | 6 | 22 | 0.893 | ns |
| shaved | 6 | 25 | 1 | ns |
| shaved | 6 | 28 | 0.055 | ns |
| shaved | 6 | 30 | 1.35e-06 | **** |
| shaved | 10 | 14 | 1 | ns |
| shaved | 10 | 18 | 1 | ns |
| shaved | 10 | 22 | 1 | ns |
| shaved | 10 | 25 | 1 | ns |
| shaved | 10 | 28 | 0.19 | ns |
| shaved | 10 | 30 | 3.72e-06 | **** |
| shaved | 14 | 18 | 1 | ns |
| shaved | 14 | 22 | 1 | ns |
| shaved | 14 | 25 | 1 | ns |
| shaved | 14 | 28 | 0.286 | ns |
| shaved | 14 | 30 | 1.41e-05 | **** |
| shaved | 18 | 22 | 1 | ns |
| shaved | 18 | 25 | 1 | ns |
| shaved | 18 | 28 | 1 | ns |
| shaved | 18 | 30 | 3.3e-04 | *** |
| shaved | 22 | 25 | 1 | ns |
| shaved | 22 | 28 | 1 | ns |
| shaved | 22 | 30 | 0.001 | ** |
| shaved | 25 | 28 | 0.428 | ns |
| shaved | 25 | 30 | 0.001 | ** |
| shaved | 28 | 30 | 0.001 | ** |

| $T_a$ | mouse group | Mean RER | SD | N |
|---|---|---|---|---|
| 6 | furry | 0.91 | 0.013 | 9 |
| 6 | shaved | 0.905 | 0.014 | 9 |
| 10 | furry | 0.924 | 0.033 | 9 |

| | | | | |
|---|---|---|---|---|
| 10 | shaved | 0.932 | 0.028 | 9 |
| 14 | furry | 0.911 | 0.04 | 9 |
| 14 | shaved | 0.927 | 0.047 | 9 |
| 18 | furry | 0.918 | 0.059 | 9 |
| 18 | shaved | 0.952 | 0.023 | 9 |
| 22 | furry | 0.951 | 0.032 | 9 |
| 22 | shaved | 0.949 | 0.04 | 9 |
| 25 | furry | 0.962 | 0.056 | 9 |
| 25 | shaved | 0.892 | 0.082 | 9 |
| 28 | furry | 0.965 | 0.028 | 9 |
| 28 | shaved | 1.004 | 0.062 | 9 |
| 30 | furry | 1.051 | 0.064 | 9 |
| 30 | shaved | 1.142 | 0.041 | 9 |

**Table S4. Role of insulation in metabolism and thermoregulation of naked mole-rats. Related to Figure 6.**

**Effect of insulation across tested $T_a$ groups on NMR $T_b$**
**Two-way repeated measures ANOVA (type III tests)**

| Effect | DFn | DFd | F | p | ges |
|---|---|---|---|---|---|
| $T_a$ | 4 | 16 | 405.644 | 7.37e-16 | 0.96 |
| insulation | 1 | 4 | 38.894 | 0.003 | 0.55 |
| $T_a$ : insulation | 4 | 16 | 9.763 | 0.00034 | 0.41 |

**Effect of insulation at each $T_a$ group**
**Pairwise comparisons of mean $T_b$ between insulated and control NMRs (paired t-test, Bonferroni adj.)**

| $T_a$ | group 1 | group 2 | p.adj | p.adj.signif |
|---|---|---|---|---|
| 18 | control | insulated | 0.003 | ** |
| 22 | control | insulated | 0.09 | ns |
| 25 | control | insulated | 0.002 | ** |
| 28 | control | insulated | 0.427 | ns |
| 32 | control | insulated | 0.028 | * |

**Effect of $T_a$ on NMR $T_b$ – insulated group**
**Pairwise comparisons of mean $T_b$ in insulated NMRs (paired t-test, Bonferroni adj.)**

| insulation | group 1 | group 2 | p.adj | p.adj.signif |
|---|---|---|---|---|
| insulated | 18 | 22 | 0.169 | ns |
| insulated | 18 | 25 | 0.003 | ** |
| insulated | 18 | 28 | 0.001 | ** |
| insulated | 18 | 32 | 0.0004 | *** |
| insulated | 22 | 25 | 0.008 | ** |
| insulated | 22 | 28 | 0.019 | * |
| insulated | 22 | 32 | 0.002 | ** |
| insulated | 25 | 28 | 0.213 | ns |
| insulated | 25 | 32 | 0.00047 | *** |
| insulated | 28 | 32 | 0.066 | ns |

| $T_a$ | NMR group | Mean $T_b$ | SD | N |
|---|---|---|---|---|
| 18 | control | 21.3 | 1.04 | 5 |
| 18 | insulated | 25.2 | 1.11 | 5 |
| 22 | control | 26 | 0.43 | 5 |
| 22 | insulated | 27.3 | 1.22 | 5 |
| 25 | control | 28.5 | 0.65 | 5 |
| 25 | insulated | 30.4 | 0.67 | 5 |
| 28 | control | 31.1 | 0.8 | 5 |
| 28 | insulated | 31.6 | 1.05 | 5 |
| 32 | control | 33.1 | 0.38 | 5 |
| 32 | insulated | 33.7 | 0.32 | 5 |

**Effect of insulation across tested $T_a$ groups on NMR EE**
**Two-way repeated measures ANOVA (type III tests)**

| Effect | DFn | DFd | F | p | ges |
|---|---|---|---|---|---|
| $T_a$ | 4 | 16 | 17.238 | 0.0000119 | 0.60 |
| insulation | 1 | 4 | 126.507 | 0.000356 | 0.58 |
| $T_a$ : insulation | 4 | 16 | 6.287 | 0.003 | 0.31 |

**Effect of insulation at each $T_a$ group**
**Pairwise comparisons of mean EE between insulated and control NMRs (paired t-test, Bonferroni adj.)**

| $T_a$ | group 1 | group 2 | p.adj | p.adj.signif |
|---|---|---|---|---|
| 18 | control | insulated | 0.042 | * |
| 22 | control | insulated | 0.00037 | *** |

| | | | | |
|---|---|---|---|---|
| 25 | control | insulated | 0.018 | ns |
| 28 | control | insulated | 0.427 | * |
| 32 | control | insulated | 0.103 | ns |

**Effect of $T_a$ on NMR EE**
**One-way repeated measures ANOVA (type III tests)**

| insulation | Effect | F | p | ges |
|---|---|---|---|---|
| control | $T_a$ | 11.5 | 0.00013 | 0.69 |
| insulated | $T_a$ | 16.5 | 0.000015 | 0.579 |

**Pairwise comparisons of mean EE in insulated and control NMRs (paired t-test, Bonferroni adj.)**

| insulation | group 1 | group 2 | p.adj | p.adj.signif |
|---|---|---|---|---|
| control | 18 | 22 | 1 | ns |
| control | 18 | 25 | 1 | ns |
| control | 18 | 28 | 1 | ns |
| control | 18 | 32 | 0.192 | ns |
| control | 22 | 25 | 0.399 | ns |
| control | 22 | 28 | 0.797 | ns |
| control | 22 | 32 | 0.011 | * |
| control | 25 | 28 | 1 | ns |
| control | 25 | 32 | 0.001 | ** |
| control | 28 | 32 | 0.023 | * |
| insulated | 18 | 22 | 0.679 | ns |
| insulated | 18 | 25 | 1 | ns |
| insulated | 18 | 28 | 0.797 | ns |
| insulated | 18 | 32 | 0.018 | * |
| insulated | 22 | 25 | 0.258 | ns |
| insulated | 22 | 28 | 1 | ns |
| insulated | 22 | 32 | 0.009 | ** |
| insulated | 25 | 28 | 0.048 | * |
| insulated | 25 | 32 | 0.034 | * |
| insulated | 28 | 32 | 0.495 | ns |

| $T_a$ | NMR group | Mean EE | SD | N |
|---|---|---|---|---|
| 18 | control | 0.275 | 0.063 | 5 |
| 18 | insulated | 0.201 | 0.026 | 5 |
| 22 | control | 0.339 | 0.044 | 5 |
| 22 | insulated | 0.173 | 0.021 | 5 |
| 25 | control | 0.261 | 0.038 | 5 |
| 25 | insulated | 0.209 | 0.042 | 5 |
| 28 | control | 0.264 | 0.058 | 5 |
| 28 | insulated | 0.161 | 0.043 | 5 |
| 32 | control | 0.144 | 0.023 | 5 |
| 32 | insulated | 0.118 | 0.011 | 5 |

**F-statistics on RER means of NMRs across tested $T_a$ groups**
**Two-way repeated measures ANOVA (type III tests)**

| Effect | DFn | DFd | F | p | ges |
|---|---|---|---|---|---|
| $T_a$ | 4 | 16 | 11.204 | 0.000158 | 0.64 |
| insulation | 1 | 4 | 170.013 | 0.0002 | 0.54 |
| $T_a$ : insulation | 4 | 16 | 12.06 | 0.000104 | 0.4 |

**Effect of insulation at each $T_a$ group**
**Pairwise comparisons of mean $T_b$ between insulated and control NMRs (paired t-test, Bonferroni adj.)**

| $T_a$ | group 1 | group 2 | p.adj | p.adj.signif |
|---|---|---|---|---|
| 18 | control | insulated | 0.003 | ** |

| | | | | |
|---|---|---|---|---|
| 22 | control | insulated | 0.09 | ns |
| 25 | control | insulated | 0.002 | ** |
| 28 | control | insulated | 0.427 | ns |
| 32 | control | insulated | 0.028 | * |

**Effect of $T_a$ on NMR $T_b$ – insulated group**
**Pairwise comparisons of mean $T_b$ in insulated NMRs (paired t-test, Bonferroni adj.)**

| insulation | group 1 | group 2 | p.adj | p.adj.signif |
|---|---|---|---|---|
| insulated | 18 | 22 | 0.169 | ns |
| insulated | 18 | 25 | 0.003 | ** |
| insulated | 18 | 28 | 0.001 | ** |
| insulated | 18 | 32 | 0.0004 | *** |
| insulated | 22 | 25 | 0.008 | ** |
| insulated | 22 | 28 | 0.019 | * |
| insulated | 22 | 32 | 0.002 | ** |
| insulated | 25 | 28 | 0.213 | ns |
| insulated | 25 | 32 | 0.00047 | *** |
| insulated | 28 | 32 | 0.066 | ns |

| $T_a$ | NMR group | Mean $T_b$ | SD | N |
|---|---|---|---|---|
| 18 | control | 21.3 | 1.04 | 5 |
| 18 | insulated | 25.2 | 1.11 | 5 |
| 22 | control | 26 | 0.43 | 5 |
| 22 | insulated | 27.3 | 1.22 | 5 |
| 25 | control | 28.5 | 0.65 | 5 |
| 25 | insulated | 30.4 | 0.67 | 5 |
| 28 | control | 31.1 | 0.8 | 5 |
| 28 | insulated | 31.6 | 1.05 | 5 |
| 32 | control | 33.1 | 0.38 | 5 |
| 32 | insulated | 33.7 | 0.32 | 5 |

**Effect of insulation across tested $T_a$ groups on NMR RER**
**Two-way repeated measures ANOVA (type III tests)**

| Effect | DFn | DFd | F | p | ges |
|---|---|---|---|---|---|
| $T_a$ | 4 | 16 | 11.204 | 0.000158 | 0.64 |
| insulation | 1 | 4 | 170.013 | 0.0002 | 0.54 |
| $T_a$ : insulation | 4 | 16 | 12.06 | 0.0001 | 0.4 |

**Effect of insulation at each $T_a$ group**
**Pairwise comparisons of mean RER between insulated and control NMRs (paired t-test, Bonferroni adj.)**

| $T_a$ | group 1 | group 2 | p.adj | p.adj.signif |
|---|---|---|---|---|
| 18 | control | insulated | 0.956 | ns |
| 22 | control | insulated | 0.375 | ns |
| 25 | control | insulated | 0.000038 | **** |
| 28 | control | insulated | 0.009 | ** |
| 32 | control | insulated | 0.004 | ** |

**Effect of $T_a$ on NMR RER**
**Pairwise comparisons of mean EE in insulated and control NMRs (paired t-test, Bonferroni adj.)**

| insulation | group 1 | group 2 | p.adj | p.adj.signif |
|---|---|---|---|---|
| control | 18 | 22 | 0.043 | * |
| control | 18 | 25 | 1 | ns |
| control | 18 | 28 | 1 | ns |
| control | 18 | 32 | 0.637 | ns |
| control | 22 | 25 | 0.282 | ns |
| control | 22 | 28 | 1 | ns |
| control | 22 | 32 | 1 | ns |

| | | | | |
|---|---|---|---|---|
| control | 25 | 28 | 1 | ns |
| control | 25 | 32 | 0.56 | ns |
| control | 28 | 32 | 0.857 | ns |
| insulated | 18 | 22 | 0.247 | ns |
| insulated | 18 | 25 | 0.019 | * |
| insulated | 18 | 28 | 0.003 | ** |
| insulated | 18 | 32 | 0.00099 | *** |
| insulated | 22 | 25 | 0.078 | ns |
| insulated | 22 | 28 | 0.295 | ns |
| insulated | 22 | 32 | 0.048 | * |
| insulated | 25 | 28 | 1 | ns |
| insulated | 25 | 32 | 0.098 | ns |
| insulated | 28 | 32 | 0.093 | ns |

| **$T_a$** | **NMR group** | **Mean RER** | **SD** | **N** |
|---|---|---|---|---|
| 18 | control | 0.839 | 0.023 | 5 |
| 18 | insulated | 0.839 | 0.021 | 5 |
| 22 | control | 0.898 | 0.029 | 5 |
| 22 | insulated | 0.913 | 0.031 | 5 |
| 25 | control | 0.838 | 0.025 | 5 |
| 25 | insulated | 0.964 | 0.029 | 5 |
| 28 | control | 0.821 | 0.079 | 5 |
| 28 | insulated | 0.973 | 0.019 | 5 |
| 32 | control | 0.936 | 0.081 | 5 |
| 32 | insulated | 1.066 | 0.036 | 5 |

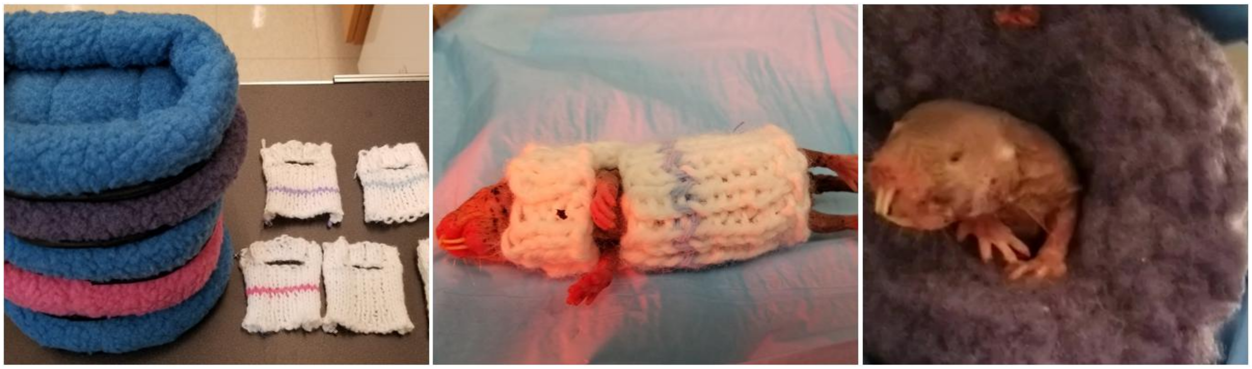

**Fig. S4. Insulation methods for naked mole-rats (NMRs) in thermoregulation studies.**
Left panel: Custom-made wool sweaters (right) and fleece beds (left) used as insulation methods for NMRs. The sweaters were the initial attempt at providing constant insulation, while the fleece beds were ultimately chosen as the preferred method. Middle panel: An NMR wearing a custom-made wool sweater. This method was discontinued due to signs of distress in the animals and their attempts to remove the garments. Right panel: An NMR voluntarily using a fleece bed insulation enclosure. This method was chosen for the study as it allowed for natural burrowing behavior and minimized stress on the animals.
This figure illustrates the evolution of our insulation methodology, from the initially attempted constant insulation approach to the final voluntary use design that better accommodated NMR behavior and well-being.

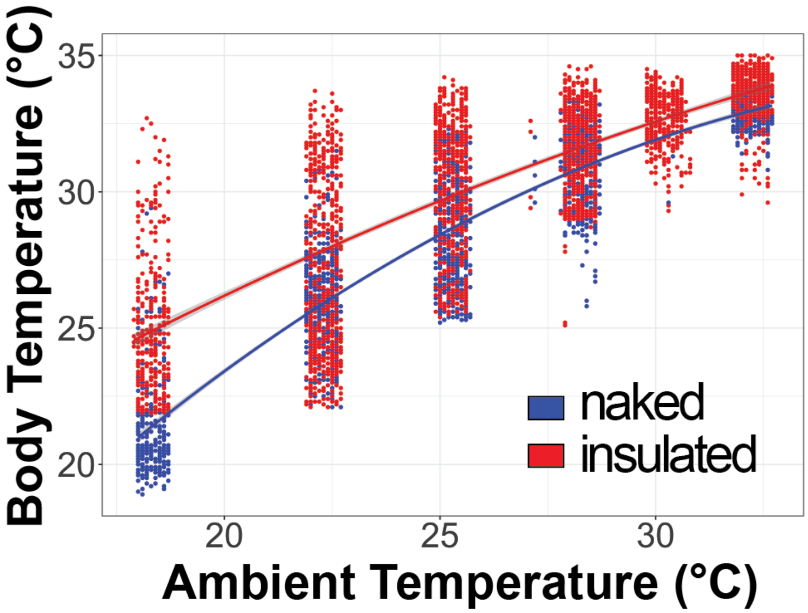

**Fig. S5. Variability in core body temperature measurements of insulated naked mole-rats (NMRs) due to voluntary use of fleece bed enclosures.** Each point represents the hourly average core body temperature of one NMR (n=5) housed at the specified ambient temperature (Ta). At ambient temperatures of 18 and 22 °C, insulated NMRs demonstrate an increase in body temperature by up to 10 °C at maximum values. Despite this variability, insulated NMRs maintained higher average body temperatures compared to non-insulated controls (see Fig. 6A).

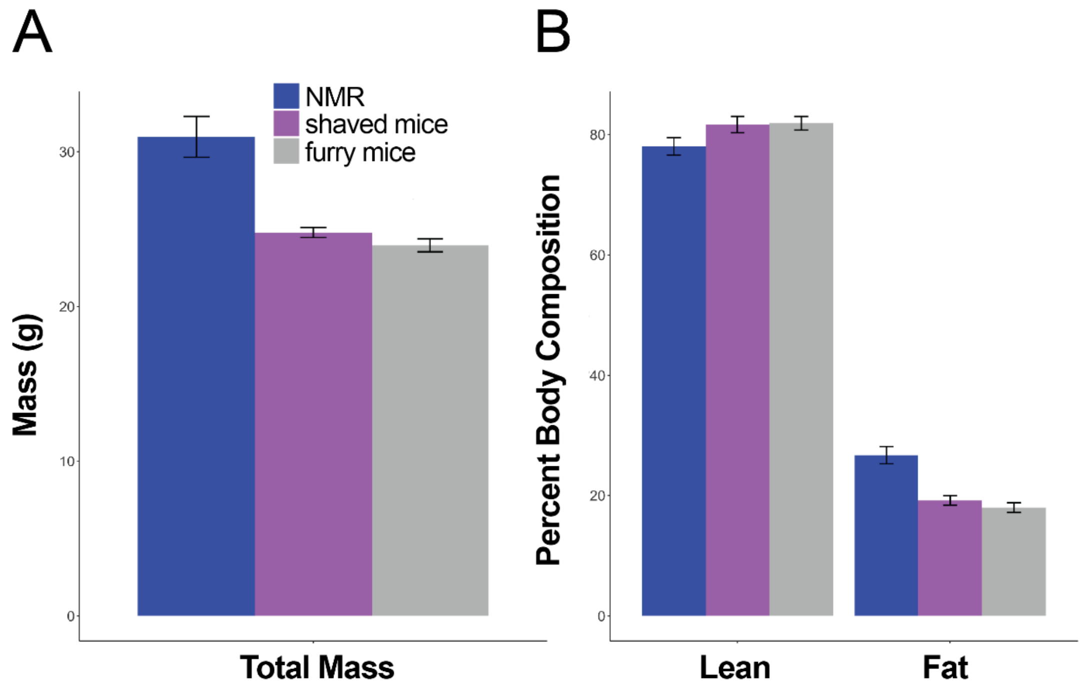

**Fig. S6. Body mass and composition of NMRs and mice as measured by nuclear magnetic resonance**. (*A*) Total body mass of the three groups of animals used for the temperature challenge experiments. (*B*) Percent body composition.
